# IDEAL-Age: an interpretable deep learning framework for single-cell resolution profiling of immunological aging

**DOI:** 10.64898/2025.12.25.696528

**Authors:** Yin Xu, Zhengchao Luo, Kai He, Feifan Zhang, Yawei Zhang, Jinzhuo Wang, Han Wen, Yongge Li, Dali Han

## Abstract

Immunosenescence increases susceptibility to infection and reduces vaccine responsiveness, yet bulk transcriptomic clocks obscure the cellular heterogeneity underlying this process. Here, we present IDEAL-Age, an interpretable deep learning framework that operates directly on single-cell PBMC transcriptomes. Benchmarking against 31 methods across independent cohorts demonstrates superior predictive performance. The framework’s interpretability uncovers linear and non-linear transcriptomic dynamics that reveal phase-specific physiological transitions, and identifies pro-youthful or pro-aging cellular contributions. Application to systemic lupus erythematosus (SLE) reveals accelerated immunological aging driven by interferon-associated monocyte shifts. IDEAL-Age establishes a high-resolution computational framework for deciphering systemic immune aging.

## Background

Aging is a time-dependent functional decline that affects nearly all organisms and is the primary risk factor for major human pathologies, including cancer, cardiovascular disorders, and neurodegenerative diseases [1, 2]. Furthermore, conditions such as HIV infection, type 2 diabetes, and systemic lupus erythematosus (SLE) are also associated with phenotypes that resemble accelerated aging [3-5]. At the cellular level, the immune system is among the first to deteriorate, a process termed immunosenescence, characterized by thymic involution, accumulation of memory/exhausted T cells, reduced T-cell receptor (TCR) diversity, and a chronic low-grade inflammatory state known as inflammaging [6, 7]. Because immune dysfunction both drives and mirrors systemic aging, quantitative assessment of immunosenescence is now viewed as a central pillar for understanding organismal aging and for designing interventions that extend healthspan [8].

Quantitative assessment of immunological age, which captures the physiological state rather than chronological time, is essential for monitoring aging dynamics and evaluating potential geroprotective interventions [9]. Over the past decade, multiple aging clocks have been devised using DNA methylation [10-12], proteomics [13, 14], metabolomics [15, 16], and imaging features [17, 18] that correlate with morbidity and mortality risk. Among these, transcriptomic aging clocks have garnered attention owing to the dynamic and responsive nature of gene expression profiles, which reflect changes in environment, metabolism, and immune status, offering real-time snapshots of physiological aging processes that some epigenetic clocks may not fully capture [19, 20]. Peripheral blood mononuclear cells (PBMCs) are an ideal surrogate tissue for such models: they are easily accessible, comprise the major players of adaptive and innate immunity, and their transcriptional programs are highly sensitive to immunosenescence [21, 22].

Current PBMC-based transcriptomic clocks predict chronological age with a mean absolute error (MAE) of 5–8 years and can detect age acceleration in cohorts with autoimmune diseases, cancer or chronic viral infections [17, 19, 22]. However, these models rely on bulk RNA-seq signals averaged across millions of cells, thereby masking the considerable heterogeneity among immune sub-populations that underlies immunosenescence. Single-cell RNA-sequencing (scRNA-seq) has revealed that only specific subsets, such as CD8^+^ CD28^-^ CD57^+^ senescent T cells or NKG2C^+^ adaptive NK cells, expand with age, whereas naïve CD4^+^ T cells and memory B cells decline [23, 24]. Bulk clocks cannot attribute age-associated expression changes to particular cell types, nor can they dissect cell-type communication networks (e.g., IFN-γ–sensing or IL-10–mediated suppression) that propagate senescence phenotypes across the immune ecosystem [25]. Consequently, existing transcriptomic age predictors fall short of linking an individual’s global aging trajectory to the aging state of each immune cell population and their interactive circuitry—knowledge that is critical for precision immune-rejuvenation strategies.

Recent attempts to leverage single-cell transcriptomics for age prediction have primarily taken a “pseudo-bulk” approach—aggregating single-cell data from individuals to approximate bulk profiles, focusing on specific cell subsets independently [25, 26], or utilizing cell-type composition characteristics to construct clocks [24]. While this enhances resolution compared to bulk RNA-seq, these methods still do not fully exploit the rich granularity of individual cell-level information and overlook the substantial transcriptional heterogeneity and functional diversity that exists within defined cell subsets. Moreover, many models lack robust interpretability at both gene and cell-type levels, limiting biological insight and clinical translatability.

To address the limitations of conventional bulk and pseudo-bulk approaches, which obscure cellular heterogeneity, we developed IDEAL-Age (Interpretable Deep Ensemble with Attention for immunoLogical Age prediction), a novel interpretable deep learning framework designed for single-cell resolution immunological age prediction from PBMC single-cell transcriptomes. Unlike previous methodologies, IDEAL-Age provides multi-scale interpretability at both the gene and individual cell levels, enabling the analysis of cellular contributions to immunological aging. Benchmarking against 31 methods across large-scale, independent cohorts (∼20 million cells) demonstrated its superior predictive accuracy and robust cross-cohort generalizability. Our results reveal conserved aging signatures alongside non-linear transcriptomic dynamics, indicating phase-specific physiological transitions that temporally align with established proteomic aging waves, while identifying pro-youthful and pro-aging cellular contributions. Finally, application to systemic lupus erythematosus (SLE) patient data reveals an accelerated immune aging phenotype driven by interferon-associated monocyte shifts, demonstrating the framework’s translational utility. Together, these findings establish IDEAL-Age as a powerful high-resolution computational framework for the precise quantification and mechanistic dissection of systemic immunological aging.

## Results

### Overview of IDEAL-Age

To fully utilize the high-resolution information embedded in single-cell transcriptomes, we developed IDEAL-Age (Interpretable Deep Ensemble with Attention for immunoLogical Age prediction), a novel framework for donor-level age prediction (Fig. 1a). Grounded in the mathematical principles of permutation-invariant architectures, the model treats each donor as an unordered collection of cells, which allows it to learn predictive signals while respecting the inherent variability in cell number and composition across individuals. This design preserves the biological resolution of single-cell data and enables a direct link between organism-level aging and molecular states at the cell level.

**Fig. 1.**
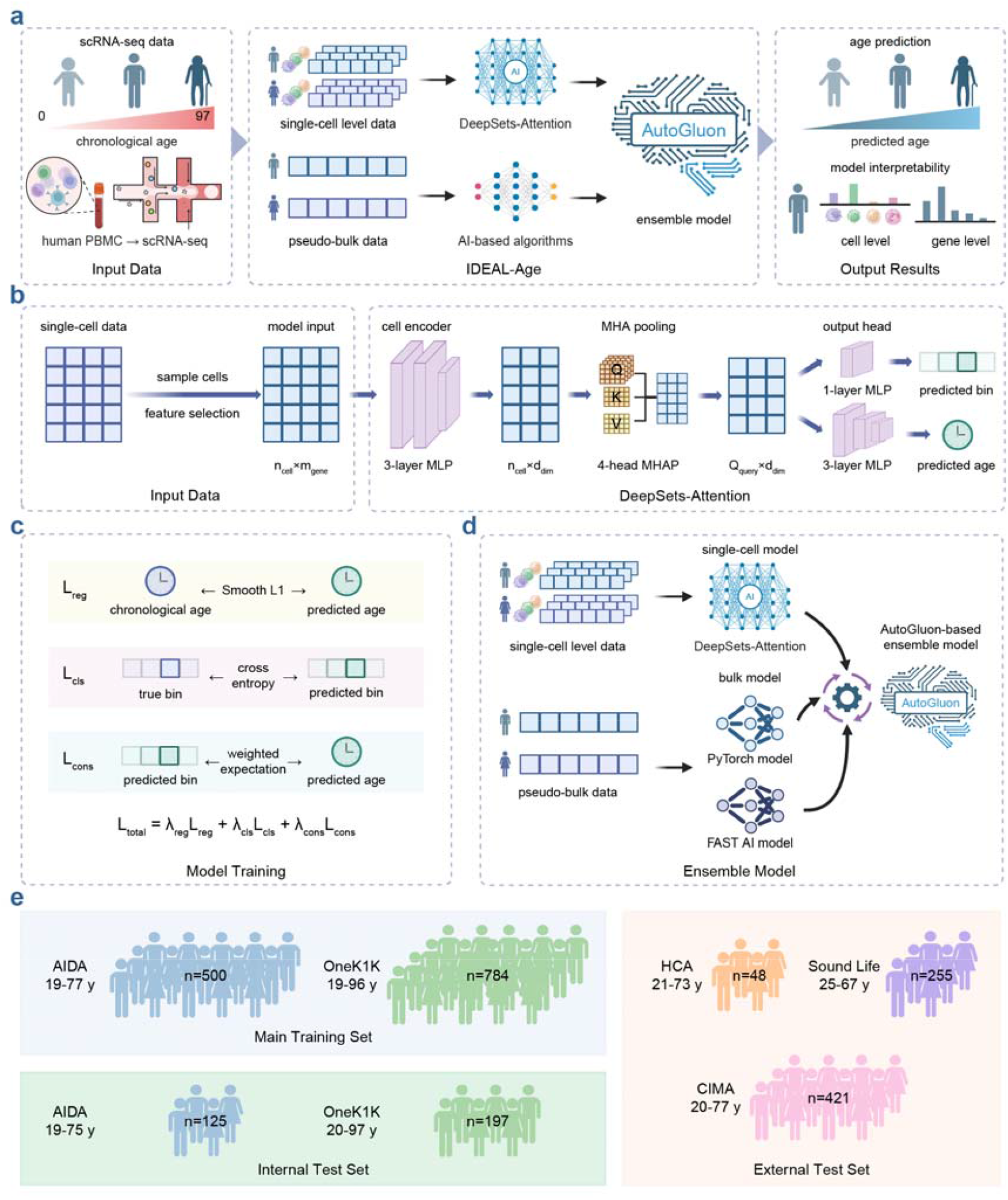
The illustration of the IDEAL-Age model. **a** Graphic overview of the IDEAL-Age model. **b** Graphic overview of the single-cell level DeepSets-Attention model. **c** Illustration of the loss function for model training. **d** Diagram of the ensemble framework. **e** Summary of the cohort information.

The single-cell level model, assigned as DeepSets-Attention, comprises three sequential modules that together process, integrate, and interpret the granular single-cell information into a donor-level age prediction (Fig. 1b).

First, a cell-level encoder represents transcriptional state of each immune cell. Each cell is first mapped to a lower-dimensional latent representation using a shared multi-layer perceptron (MLP). This encoder learns subtle, non-linear transcriptomic states while maintaining biological consistency across donors. Cells that arise in aging contexts, such as rare dysfunctional T cells or activated myeloid populations, are captured as distinct points in this latent space before any information is aggregated.

Second, a multi-head attention pooling (MHAP) layer learns how aging-relevant subsets shape donor-level phenotypes. Unlike traditional methods that use simple mean or max pooling, which treat all cells as equally informative, we implemented a multi-head attention pooling module that performs adaptive aggregation across heterogeneous cell states. Recognizing that immunosenescence is a multifaceted process driven by distinct cell subsets, this module learns a small set of trainable query vectors that probe the cellular landscape of each donor. Each query can be interpreted as a computational analogue of a biological motif, sensitized to a specific axis of immunosenescence. These queries allow the model to adaptively weigh and aggregate information from biologically relevant cells while filtering out technical noise, effectively learning which cell states are most predictive of aging without prior manual selection.

Third, a donor-level prediction head models continuous and categorical aspects of immune aging. The aggregated cell representations are finally processed by a prediction head to estimate the immunological age. To enhance generalization and enforce biological consistency, we employed a multi-task loss function that simultaneously optimizes for precise age regression (using the Smooth L1 regression loss) to capture the continuous nature of aging and broad age-group classification, which encourages the separation of broad physiological categories such as young, middle-aged, and older donors (Fig. 1c).

To ensure robustness and predictive accuracy, the donor-level prediction is constructed as an ensemble using the AutoGluon framework [27] (Fig. 1d). This strategy integrates the custom DeepSets-Attention networks with diverse baseline models trained on pseudo-bulk features, optimizing performance through stacked ensemble and capturing complementary biological signals. This architecture not only achieves superior predictive accuracy but, crucially, establishes a direct computational link between donor age and the transcriptional states of individual cells, serving as the foundation for the cell-level and gene-level interpretability analyses presented below.

To refine the model and enhance its biological interpretability, we performed a targeted feature selection procedure. The final set of features was constituted as the union of three distinct gene categories: (1) established cell-type marker genes, (2) previously reported aging-related genes from the literature, and (3) data-driven age-related genes identified via a mutual information criterion with age as the target variable (Fig. S1). Detailed parameter settings, optimization strategies for model training, and the complete implementation process of ensemble learning are provided in the Methods section.

To construct a robust training cohort, we integrated single-cell transcriptomic data from two large-scale studies: the Asian Immune Diversity Atlas (AIDA) [28] and OneK1K [29] (Fig. 1e, Fig. S2a and Table S1–S2). Specifically, 80% of donors from each cohort were randomly assigned to the training set, encompassing 500 donors (aged 19–77) from AIDA and 784 donors (aged 19–96) from OneK1K. The remaining 20% of donors from both AIDA and OneK1K served as internal test sets to evaluate the model’s performance and generalizability.

### IDEAL-Age robustly predicts biological immune age across diverse datasets

Next, we systematically evaluated the performance of IDEAL-Age by comparing its predicted immunological age with chronological ages. We first utilized internal test sets comprising 125 donors randomly selected from the AIDA dataset and 197 donors from the OneK1K dataset. Considering that many existing aging clocks face performance degradation when applied to independent datasets, we further assessed the cross-dataset generalizability of IDEAL-Age using three external test sets: the Human Cell Atlas (HCA) dataset [30], which included 48 donors; the Sound Life Project [31], consisting of 255 longitudinal samples from 96 donors (aged 25–67); and the Chinese Immune Multi-Omics Atlas (CIMA) [32], encompassing 421 donors (aged 20–77) (Fig. 1e, Fig. S2a and Table S1–S2). To minimize the confounding effects of pathological states on biological age estimation, only samples from clinically healthy individuals across these cohorts were retained for downstream analysis.

We then conducted systematic benchmarking of IDEAL-Age against published senescence evaluation tools and transcriptome-based aging clocks using five scRNA-seq datasets (two internal test sets AIDA and OneK1K, and three external cohorts HCA, Sound Life, and CIMA) derived from healthy human PBMCs. A total of 31 methods were included and categorized into four groups: 13 classical gene markers for senescent cells (referred to as “Aging Marker”, e.g., *CDKN2A*), 8 widely used senescence-associated gene sets (referred to as “Aging Set”: SASP [33], SenMayo [34], CellAge [35], SigRS [36], SenUp [37], AgingAtlas [38], ASIG [39], GenAge [40]), 3 senescence scoring methods (referred to as “Aging Score”: hUSI [41] SENCAN [42], SENCID [43]), and 7 aging clocks trained on scRNA-seq data (referred to as “Aging Clock”: composition-based clock PLSR (cell-composition) [24], siAge [26], and 5 models of scImmuAging framework [25]).

To evaluate the prediction accuracy, we calculated the Pearson correlation coefficient (PCC) and mean absolute error (MAE) between predicted immunological age and chronological age. IDEAL-Age maintained stable predictive performance across the training sets (PCC = 0.95, MAE = 3.91), internal test sets (PCC = 0.878, MAE = 6.91), and external test sets (PCC = 0.819, MAE = 7.81) (Fig. 2a, Fig. S3a-c, Table S3).

**Fig. 2.**
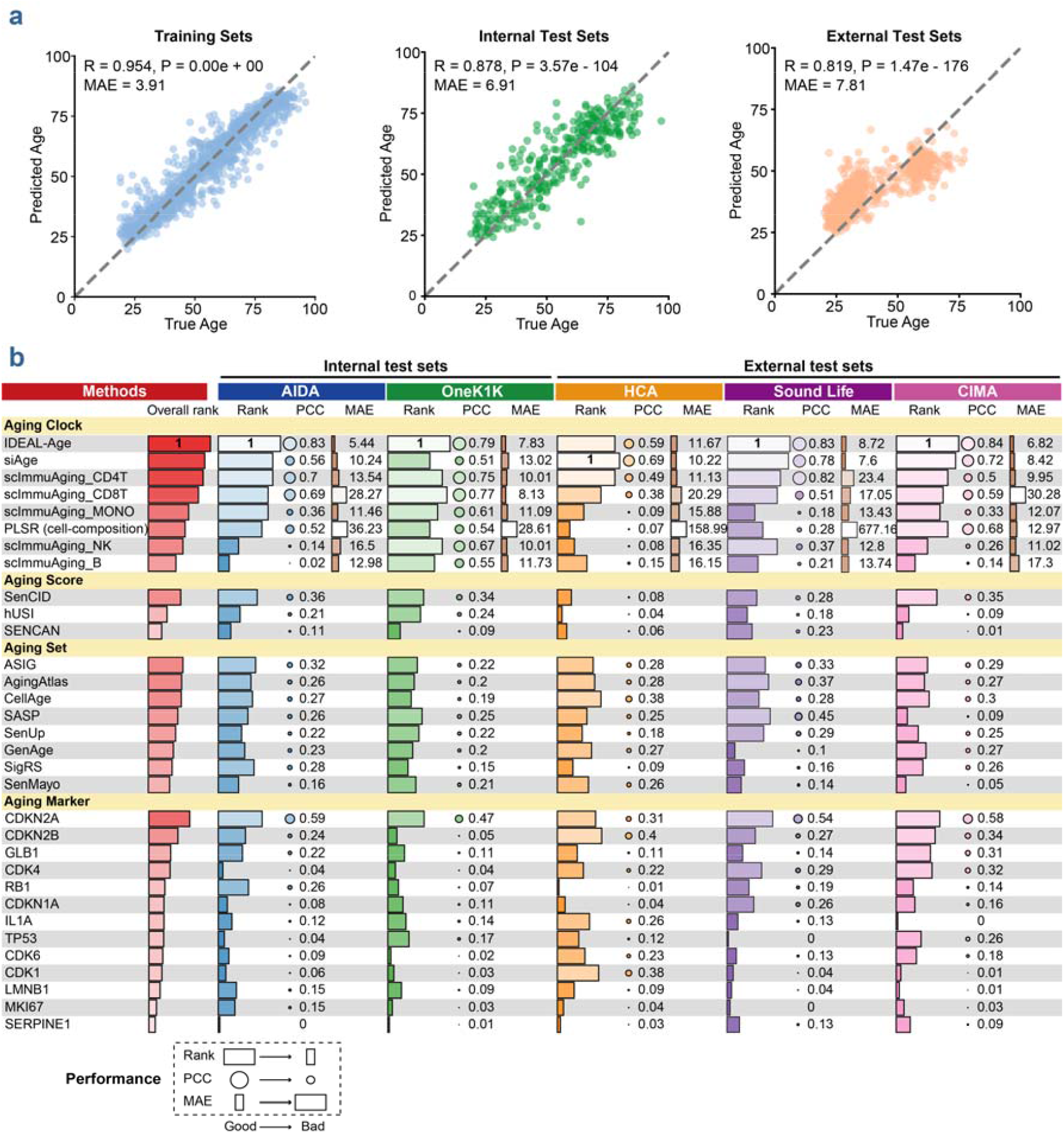
Evaluations on internal and external datasets demonstrate the accuracy and robustness of IDEAL-Age aging clock. **a** Scatter plot showing the predicted age versus chronological age across training, internal test and external test datasets of IDEAL-Age. **b** Heatmap summary of predictive performance for various aging models (rows) evaluated on five independent datasets (columns), comprising two internal test sets (AIDA and OneK1K) and three external test sets (HCA, Sound Life, and CIMA). Models are categorized by their underlying evaluation strategies (Aging Clock, Aging Score, Aging Set, and Aging Marker). The leftmost red bars display the overall rank of each method across all datasets. For each specific dataset, predictive performance is evaluated using three metrics: dataset-specific performance rank, PCC, and MAE. The legend at the bottom denotes the relationship between geometric properties and performance quality (Good to Bad).

In the comparative analysis, IDEAL-Age achieved the highest overall rank compared to 31 evaluated methods (Fig. 2b). Specifically, IDEAL-Age ranked first in both internal validation cohorts (AIDA and OneK1K) and two large-scale external cohorts (Sound Life and CIMA), and ranked second in the external HCA dataset. Notably, despite the inherent technical shifts present in out-of-distribution data, IDEAL-Age maintained highly competitive PCC and MAE values relative to existing models (Fig. 2b). This consistent performance underscores the framework’s exceptional structural robustness and its capacity for reliable biological pattern recognition across independent cross-cohort environments.

### Age-stratified evaluation and targeted boundary fine-tuning

Furthermore, we investigated the models’ predictive performance across distinct age distributions. The baseline model, trained predominantly on adult cohorts (19–77 years), naturally underrepresented the non-linear transcriptomic dynamics characteristic of pediatric development (<19 years) and advanced age (>77 years), resulting in boundary biases such as a pediatric “floor effect” (Fig. S4, Table S2 and S4). To address this, we implemented a targeted fine-tuning strategy utilizing supplementary samples from 2 tuning cohorts containing extreme-age donors (siAge study with 60 donors aged 0–90 [26] and the SC2018 dataset with 7 centenarians [30], Fig. S2, Table S1–S2). To determine the optimal bias-mitigation strategy, we evaluated multiple fine-tuning configurations using these supplementary data. The model fine-tuned exclusively on the extreme-age subset of the siAge dataset—designated as IDEAL-Age (tuning)—yielded the most robust empirical improvements (Fig. S4 and S5, Table S5–S7).

Specifically, by incorporating the pediatric samples from the siAge dataset, the model eliminated the previous ‘floor effect’ in subjects under 19, reducing the MAE from 17.64 to 1.52 and improving the PCC from 0.58 to 0.95 (Fig. S4 and S5). We must transparently note that due to the current scarcity of independent pediatric PBMC scRNA-seq cohorts, an out-of-distribution test set is not yet available for the <19 age group. Consequently, this specific evaluation utilizes the pediatric subset of the siAge dataset—which also serves as the training set for both the baseline siAge model and our IDEAL-Age (tuning) model. While these metrics therefore reflect in-sample fitting performance rather than cross-cohort generalizability, this improvement suggests that our framework’s architecture possesses the necessary capacity to accurately capture early-life aging trajectories when provided with adequate data representation.

Furthermore, at the extreme aging boundary (>77 years), the model achieved an improved linear correlation (PCC increasing from 0.17 to 0.35) while maintaining a stable MAE (9.32) on strictly separated test sets (Fig. S4 and S5). Crucially, these boundary enhancements did not induce catastrophic forgetting, as the model successfully maintained its robust predictive accuracy for the central adult range (19-77 years) across the massive external test cohorts (HCA, Sound Life, and CIMA) (Fig. S4 and S5).

In summary, these comprehensive results highlight the robustness and broad applicability of IDEAL-Age across diverse datasets and across variable age distributions. By effectively leveraging single-cell resolution, our framework significantly advances the field, strengthening its potential for real-world deployment where heterogeneous data sources and out-of-training-distribution samples are commonplace.

### Ablation test of model architecture and feature selection

To quantify the specific contributions of framework components to predictive performance, we conducted an ablation analysis focusing on the model architecture and the feature selection strategy.

First, we evaluated the model architecture by comparing three configurations: a pseudo-bulk-only model, a DeepSets-Attention-only model, and the integrated IDEAL-Age model. Across the training, internal test, and external test datasets, the IDEAL-Age model achieved the highest predictive accuracy (Fig. S6 and Table S6–7). Quantitatively, the ensemble approach yielded average improvements of 2% in PCC and 3% in MAE compared to the pseudo-bulk-only baseline, and a 12% improvement in both metrics compared to the DeepSets-Attention-only model.

Although the predictive performance gain of the ensemble model over the pseudo-bulk baseline is quantitatively modest, the inclusion of the DeepSets-Attention branch enables single-cell-level analysis. While pseudo-bulk aggregation obscures cell-specific transcriptional variance, integrating the DeepSets-Attention module preserves single-cell resolution, establishing the computational foundation for the model’s interpretability framework. This structural ablation indicates that combining these biological scales improves predictive accuracy while maintaining the resolution required for downstream biological interpretation.

Second, to validate the necessity of our feature selection strategy, we conducted an ablation analysis comparing the model utilizing the selected 5,012-gene set against a baseline model trained on the unselected global gene set (Methods). On the internal test sets (AIDA, OneK1K), the 5,012-gene model achieved a PCC comparable to that of the global gene set model, albeit with a slightly lower MAE: 5.44 vs. 5.79 in AIDA; 7.83 vs. 7.92 in OneK1K (Fig. S7, Table S6–S7). However, substantial divergence occurred during cross-cohort evaluation on independent external test sets. While the global gene set model showed a moderate performance advantage on the HCA dataset, its predictive accuracy degraded on the Sound Life cohort (MAE increasing from 8.72 to 9.99). Most notably, it suffered a marked performance decline on the large-scale CIMA cohort, with the PCC dropping substantially from 0.84 to 0.47 and the MAE increasing from 6.82 to 9.97 (Fig. S7).

These benchmarking results demonstrate that in cross-cohort single-cell transcriptomics, the high-dimensional feature space of the full genome introduces substantial technical noise and platform-specific biases. Despite large cellular volumes, this unconstrained dimensionality leads to severe overfitting and compromises universal age prediction. Consequently, our guided 5,012-gene selection provides a necessary balance between enabling biological discovery and ensuring robust cross-cohort generalizability. Collectively, these ablation analyses demonstrate that both the model architecture and the feature selection are essential framework components. Together, they enable IDEAL-Age to achieve stable, cross-cohort predictive accuracy while preserving the necessary single-cell resolution for downstream biological interpretability.

### IDEAL-Age reveals conserved and dataset-specific molecular aging features

Following the benchmark evaluation, we next focused on how IDEAL-Age attributes donor-level predictions to the transcriptional signals of individual genes. Leveraging the model’s cell-resolved architecture, we quantified the contribution of each gene to the predicted immune age of each donor, enabling a direct mapping from single-cell expression patterns to donor-specific molecular features (Fig. 3a). Specifically, we utilized integrated gradients (IG) to quantify global gene importance (Methods). The interpretability interface demonstrated high robustness within paired internal training and testing sets, with 48 out of the top 50 genes consistently identified (Fig. S8a). This consistency highlights the model’s reliability in identifying key aging-associated genes that are broadly relevant. Specifically, genes such as *CD8B* and *RPS4Y1* consistently ranked among the top 10 contributors across four datasets (AIDA, OneK1K, HCA, and siAge) (Fig. S8b), and their age-related expression trajectories showed universal age-related dynamics characterized by significant fluctuations across the lifespan in all four independent cohorts (Fig. S8c-d). These findings support the model’s ability to detect conserved molecular signatures of immunological aging that extend beyond cohort-specific variations.

**Fig. 3.**
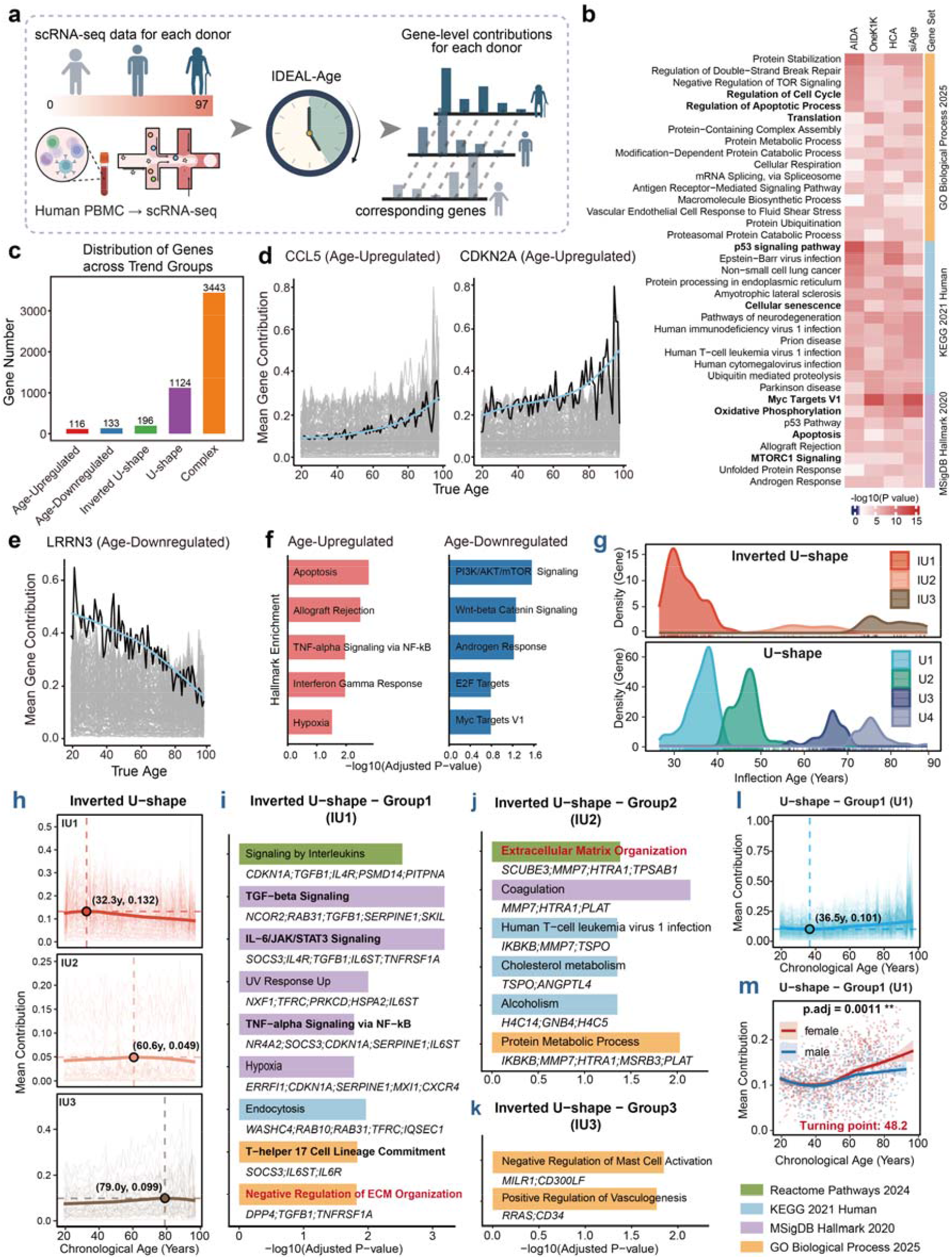
The gene-level interpretation for the aging process in 4 healthy human PBMC datasets. **a** Schematic for the gene-level interpretive interface. **b** Heatmap showing the functional enrichment analyses of the top 10% contributing genes for each dataset. *P*-values were determined by hypergeometric test. Color bars represent the gene set used for functional enrichment analyses. **c** Bar plot showing the number of genes in five trend groups based on the gene contribution. **d-e** Line plots showing example genes, (**d**) CCL5 and CDKN2A, in the age-upregulated group, and (**e**) LRRN3, in the age-downregulated group. Black lines represent the mean contribution of the highlighted genes in each age. Blue lines represent the LOWESS trends of the black lines. **f** Bar plots showing the functional enrichment terms of age-upregulated (left) and age-downregulated (right) genes in the Hallmark gene set. **g** Density plots showing the distribution patterns of inflection ages of each subgroup of inverted U-shape (upper) and U-shape (bottom) defined by *k*-means clustering. **h** Longitudinal trajectories of mean contribution across chronological age for each group with inverted U-shape. Faded thin lines represent the trajectories of individual genes, while the bold solid lines represent the LOWESS average trajectory. Dashed lines and colored circles indicate the exact group-level inflection point (peak for inverted U-shape), with the precise age and mean contribution coordinates labeled. **i-k** Pathway enrichment analysis for the inverted U-shape groups (IU1-IU3). Bar lengths represent -log_10_(Adjusted P-value) for selected significantly enriched pathways (adjusted *P*-value < 0.05). **l** Longitudinal trajectories of mean contribution across chronological age for each group with U-shape group1 (U1). **m** Sex-stratified mean contributions across chronological age for U-shape group1. Dots represent individual samples. Solid lines depict LOWESS trends with shaded areas representing 95% confidence intervals.

The distribution of gene contribution scores reveals that a small subset of genes exhibits significantly higher contribution levels (Fig. S9a). We extended our analysis by selecting the top 10% contributing genes from each dataset for functional enrichment analysis (Methods). Despite heterogeneity in the distribution of these high-contribution genes, core enriched pathways—including Myc Targets, oxidative phosphorylation, cellular senescence, apoptosis, and translation, were consistently observed across all cohorts (Fig. 3b, Fig. S9b-c). This concordance aligns well with known biological processes driving aging and immunosenescence [44-46], supporting the biological relevance of the model’s shared gene signatures.

Interestingly, when we compared gene-level interpretability across different test datasets, substantial heterogeneity emerged (Fig. S8e), indicating the model’s capacity to capture dataset-specific explanatory patterns. For example, *CLDN11* was highly influential exclusively in the HCA dataset, exhibiting clear age-associated dynamics only within this cohort (Fig. S8b, f, and g). Conversely, *NPM3* showed markedly elevated contributions and pronounced age-related expression changes primarily in the OneK1K dataset (Fig. S8b, f, and h). These observations highlight the model’s capacity for identifying context-dependent gene-age relationships, reflecting unique biological variations pertinent to immune aging in distinct cohorts.

Taken together, these results demonstrate that our model not only robustly identifies globally relevant aging signatures, but also adapts to dataset-specific variations, offering a nuanced and biologically meaningful interpretation of the complex molecular landscape underlying immune aging.

### Fine-grained donor-level gene contributions reveal biologically meaningful gene-age dynamics

Leveraging the unique donor-level interpretability interface of our IDEAL-Age model, we systematically investigated the dynamic trajectories of gene contributions with respect to donor age. We computed gene contributions averaged across donors stratified by age and categorized these trajectories into five distinct trend patterns: age-upregulated, age-downregulated, inverted U-shape, U-shape, and complex (Fig. 3c, Fig. S10) (Methods).

Interestingly, we found that genes associated with the Senescence-Associated Secretory Phenotype (SASP), including *CDKN2A* and *CCL5*, were predominantly grouped within the age-upregulated group (Fig. 3d). This suggests that in older donors, the elevated expression of these classical aging markers exerts a greater influence on age prediction by our model, consistent with previous knowledge of their roles in cellular senescence and immune aging [41, 47-49]. Conversely, genes like *LRRN3*, known to be highly expressed in naïve T cells and whose downregulation correlates with T cell aging and functional decline [15], fell into the age-downregulated group (Fig. 3e). These monotonic age-related trends highlight the model’s capability to capture both upregulated and downregulated age-associated gene expression dynamics in a fine-grained and biologically meaningful manner.

To further elucidate the biological underpinnings of these monotonic gene groups, we performed functional enrichment analysis on the genes in the age-upregulated and age-downregulated groups, respectively. The age-upregulated group showed enrichment for pathways involved in apoptosis, TNFα signaling, interferon-gamma response, and hypoxia (Fig. 3f)—processes implicated in inflammatory and stress responses during aging and immunosenescence [24, 50]. In contrast, the age-downregulated group was significantly enriched for pathways including PI3K/AKT/mTOR signaling, Wnt/β-Catenin signaling, and Myc Targets (Fig. 3f)—hallmarks of cellular metabolism, growth, and proliferation known to decline with aging [51, 52].

Together, these findings demonstrate that our model disentangles basic and opposing gene-age dynamics at the donor level. This refined molecular aging signature captures the progressive activation of pro-senescent inflammatory pathways alongside the steady decline of metabolic and proliferative signals. However, these monotonic trends represent only a foundational layer of the aging process. To better resolve the biological complexity of immunosenescence, it is necessary to explore more intricate, non-monotonic contribution patterns that reflect stage-specific physiological transitions.

### Non-monotonic gene contribution trajectories reveal stage-specific regulatory shifts in immune aging

While monotonic trends characterize the progressive divergence of aging hallmarks, they often oversimplify the dynamic nature of immunological aging, which involves complex biological transitions. To move beyond these linear approximations, we leveraged the non-linear modeling capacity of IDEAL-Age to systematically investigate gene contribution trajectories that undergo distinct phase shifts across the lifespan (the inverted U-shape and U-shape groups). By identifying inflection ages (temporal peaks or valleys) through locally weighted scatterplot smoothing (LOWESS) and applying k-means clustering, we uncovered seven data-driven clusters: three inverted U-shape (IU1–3) and four U-shape (U1–4) groups (Fig. 3g-h, Fig. S11a-b, Methods).

The inverted U-shape groups (IU1–3) characterize biological processes that reach a functional peak before declining in model predictive reliance. Specifically, IU1 (peak ∼32.3y) is enriched in core adaptive pathways, including interleukin signaling and Th17 lineage commitment (Fig. 3h, i). The model identifies the robust state of adaptive immunity as a primary predictive feature in early adulthood, with the subsequent decline in contribution mirroring the recognized timeline of thymic involution and the exhaustion of the naïve T-cell repertoire [53, 54]. IU2 (peak ∼60.6y) shows an increased model reliance on ECM organization, coagulation, and cholesterol metabolism (Fig. 3h, j). This is consistent with a recognized mid-to-late life physiological shift toward vascular and connective tissue remodeling [55, 56]. The late-life increase in IU3 (peak ∼79.0y) is driven by genes related to vasculogenesis and the negative regulation of mast cell activation (Fig. 3h, k). This delayed increase in predictive importance highlights a late-life shift in immune homeostasis, likely reflecting the critical regulation of bone marrow-derived cells required to mitigate microvascular aging [57].

While these three groups were derived entirely from the unsupervised, data-driven optimization of our deep learning model, their computational turning points (∼32.3y, 60.6y, and 79.0y) exhibit a temporal concordance with the three major waves of human plasma proteomic aging (34y, 60y, and 78y) independently established in landmark literature [58]. Furthermore, the functional progression identified by our model—from early-adulthood adaptive immunity and TGF-beta regulation (IU1) to late-life vascular and connective tissue remodeling (IU2)—conceptually mirrors the exact biological signatures defining these macroscopic proteomic waves [58]. This biological alignment provides validation that the non-linear feature weightings extracted by IDEAL-Age are not mathematical artifacts, but effectively capture fundamental, systemic milestones of human aging.

Conversely, the U-shape groups reveal biological bottlenecks in mid-life followed by an increase in predictive importance. Specifically, U1 (nadir ∼36.5y; sex-divergence ∼48.2y) is dominated by translational machinery and mitochondrial energy metabolism (Fig. 3l, Fig. S11c, g-h). This cluster captures a mid-life metabolic shift. Notably, we observed a significant sex-specific divergence in this group’s trajectory during the fifth decade (Fig. 3m). This divergence is supported by the functional enrichment of estrogen signaling and estrogen-dependent gene expression within this cluster (Fig. S11c), aligning precisely with the timing of the physiological immune remodeling characteristic of the menopausal transition [59]. U2 (nadir ∼46.9y) is defined by a significant late-life increase in predictive importance for protein homeostasis (proteasome, unfolded protein response), interferon alpha signaling, and neurodegeneration-related pathways (Fig. S11b, d). The model’s increased reliance on these features in older age quantitatively reflects the systemic proteostatic crisis—a primary driver of both organismal aging and neurodegeneration [60].

U3 (nadir ∼66.3y) is enriched in macroautophagy, mitophagy, mTOR signaling, and longevity-regulating pathways, alongside the androgen response (Fig. S11b, e). As accumulating proteotoxic and metabolic stress impacts cellular viability in advanced age, the activation state of autophagic clearance mechanisms and mTOR nutrient sensing shifts from being a baseline maintenance process to a predictive hallmark of advanced chronological age [61, 62]. U4 (nadir ∼74.8y) exhibits a late-life increase in the predictive importance of cellular stress and innate immune pathways (Fig. S11b, f). Specifically, U4 is enriched for TP53 targets and innate immune responses (innate immune system, neutrophil degranulation). The increased model contribution of these features aligns with the concept of inflammaging—a chronic, low-grade inflammatory state characteristic of advanced age [49, 63]. Furthermore, the rising predictive value of innate immunity in the mid-70s parallels the declining importance of adaptive immune pathways (observed in the IU1 group), providing computational evidence that is consistent with the recognized systemic shift from adaptive to innate immune dominance in older populations [64].

Collectively, IDEAL-Age reveals a dynamic architecture of immunological aging beyond linear models. While monotonic trends capture progressive pro-inflammatory activation and metabolic erosion, non-monotonic trajectories—aligned with systemic proteomic waves and endocrine milestones—uncover shifting biological priorities across the lifespan. By quantifying these stage-specific and sex-stratified transitions, our model provides a refined, biologically grounded map of aging. This capacity to resolve non-linear dynamics highlights the potential of deep learning to reveal molecular hallmarks obscured in traditional linear clocks.

### Identification of pro-aging and pro-youthful cell subsets

A distinctive feature of our IDEAL-Age model is its interpretability at single-cell resolution, enabling the investigation of cell-level contributions to immunological age predictions across diverse immune cell subsets by leveraging the attention-based pooling mechanism (Fig. 4a). We first projected cell-level contribution from four single-cell datasets onto the corresponding unified UMAP embeddings annotated with harmonized cell subsets (Fig. 4b-c, Fig. S12, Methods).

**Fig. 4.**
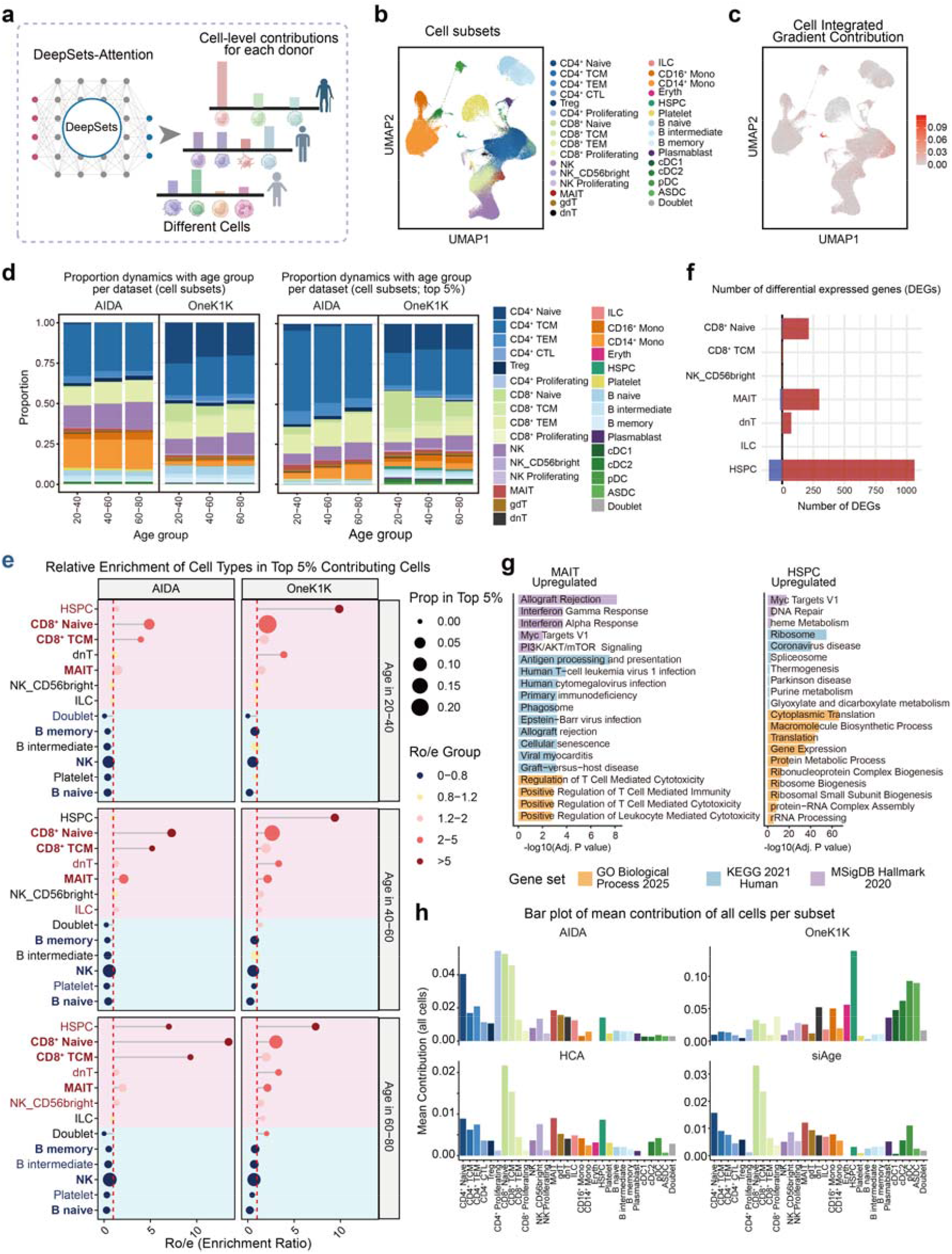
The cell-level interpretation for the aging process in 4 healthy human PBMC datasets. **a** Graphic model for the cell-level interpretability interface. **b** UMAP plot showing 31 fine-grained cell subsets identified in the integrated scRNA-seq data. **c** UMAP plot displaying the cell-level contribution scores. **d** Stacked bar plots displaying the cell subset proportions dynamics of total cells (left) and the top 5% contributing cells (right) across age groups for each dataset. **e** Lollipop plot showing the relative enrichment (Ro/e) of cell subsets across age groups and datasets. Dot size is proportional to the relative abundance of the top 5% contributing cells, and color indicating Ro/e groups. Labels in red or blue denote cell types with consistent high or low enrichment trends across datasets. Labels in bold represent cell subsets with consistent enrichment trends across age groups. The red dashed line marks the baseline Ro/e = 1. **f** Bar plot showing the number of differentially expressed genes (DEGs) for selected cell subsets between the top 5% contributing cells and the bottom 25%. Red represents the upregulated DEGs, and blue represents the downregulated. **g** Bar plots depicting functional enrichment terms of MAIT (left) and HSPC (right) upregulated genes between top 5% and bottom 25%. **h** Bar plots showing the mean contribution of each cell subset in different datasets using all cells.

To determine the biological directionality of specific cell subsets, we stratified donors from the AIDA and OneK1K test sets into accelerated, normal, and decelerated aging cohorts based on the residuals of their predicted ages (Fig. S13a-d). By evaluating shifts in relative cell proportions alongside their predictive contribution scores across these phenotypic groups (Methods), we identified distinct, directionally consistent cellular dynamics.

Specifically, NK cells, CD8^+^ T_EM_ cells, and B intermediate cells demonstrated significant proportional expansion in the accelerated cohorts across both datasets, which have been identified as robust pro-aging subsets (Fig. S13e-f). Conversely, CD4^+^ T_CM_, CD4^+^ naïve, CD8^+^ naïve, and γδ T cells exhibited consistent pro-youthful proportional shifts across both datasets (Fig. S13e-f). These computationally derived trajectories closely mirror established biological hallmarks of immunological aging, characterized by the progressive depletion of naïve T, central memory T, and γδ T cells [25, 33, 65] alongside the accumulation of effector memory T and age-associated B/NK cells [49, 66].

### Single-cell resolution interpretability uncovers intra-subset contribution heterogeneity to immunological age

To systematically evaluate how the contribution of distinct immune cell subsets varies with donor age and to ensure the robustness of observed trends, we focused on two large datasets (AIDA and OneK1K) comprising 1,446 donors aged 20–80 years. We stratified donors into three age groups (20–40, 40–60, and 60–80) and examined each dataset due to their inherent heterogeneity separately (Fig. S14a-b, Methods). The cell contribution scores follow an extremely polarized distribution, where significant contributions are confined to a very small subset of cells (Fig. S15a). Therefore, we compared the composition and cell number dynamics of all cells versus the top 5% contributing cells within each age group for each immune subset (Fig. 4d, Fig. S16).

Consistent with established immunosenescence paradigms, we observed a progressive decline in the abundance of naïve T cell subsets across increasing age groups (Fig. S14c). Conversely, CD16^+^ monocytes showed a gradual increase in representation as age advanced, reflecting known age-related myeloid expansion and pro-inflammatory shifts [67]. To better capture enrichment patterns, we calculated the relative enrichment score (Ro/e) of each cell subset within the top 5% of contributors relative to their total cell proportion for each age group (Fig. 4e, Fig. S17a). Subsets such as CD8^+^ naïve, CD8^+^ T_CM_, and MAIT cells exhibited consistent and statistically significant positive enrichment (Ro/e > 1.2) across age groups and datasets (Fig. 4e), indicating their proportionally greater involvement in driving the age-predictive signatures captured by the model. These results were consistent across different contribution cutoffs (Fig. S15b). Intriguingly, hematopoietic stem and progenitor cells (HSPCs) also showed significant positive enrichment in both the youngest (20–40 years) and oldest (60–80 years) groups (Fig. 4e), suggesting a dynamic role for this rare population in immune aging.

Altogether, seven cell subsets (CD8^+^ naïve, CD8^+^ T_CM_, MAIT, dnT, HSPC, NK_CD56bright, and ILC) demonstrated consistent compositional enrichment within the high-contribution pool (Fig. 4e). Building upon this, we sought to determine whether these contributions were uniformly distributed across each subset or driven by specific single-cell transcriptional states. Differential expression analysis revealed that only four subsets (CD8^+^ naïve, MAIT, dnT, and HSPCs) exhibited a substantial number of DEGs between their high- and low-contribution cells (Fig. 4f, Fig. S17b, Methods), highlighting significant intra-subset heterogeneity. The remaining subsets, despite being compositionally enriched, exhibited relatively homogeneous transcriptional profiles across contribution tiers. Consequently, we focused our subsequent molecular and pathway investigations exclusively on these four subsets to dissect the specific intra-subset transcriptional drivers of immunosenescence.

Subsequent functional enrichment analysis revealed that CD8^+^ naïve T and MAIT cells shared upregulated pathways including Myc targets, interferon alpha and gamma responses, and antigen processing and presentation (Fig. S17d), indicating their involvement in adaptive immunity and immune surveillance during aging [44, 66]. In contrast, double-negative T cells (dnT) displayed enrichment primarily in Myc targets and interferon responses, but also showed pronounced activation of ribosome biogenesis and translation machinery pathways, suggesting a potentially heightened cellular biosynthetic activity distinct from other subsets [68]. Similarly, HSPCs were characterized by significant enrichment of ribosome, translation, Myc target, and DNA repair pathways, reflecting their proliferative capacity and genomic maintenance critical for hematopoiesis during aging [69-71] (Fig. 4g). Notably, MAIT cells uniquely exhibited enrichment for pathways related to human cytomegalovirus (CMV) infection (Fig. 4g), linking this subset’s transcriptomic aging signature to known CMV-driven immune modulation [72].

It is worth noting that the absence of expected enrichment in certain subsets such as CD4^+^ naïve T cells in the OneK1K dataset may be attributed to the substantially high contributions from subsets like HSPCs, which could dominate the model’s attention (Fig. S17b, c). However, in two additional external datasets, CD4^+^ naïve T cells and other subsets exhibited enrichment trends consistent with established immunological knowledge (Fig. 4h), demonstrating both the robustness of our model and the biological consistency of these cell types in contributing to immune aging across independent cohorts.

Importantly, the interpretability of our model at single-cell resolution further reveals substantial heterogeneity within each immune cell subset: not all cells within a given subset contribute equally to the immunological age prediction. This capability to dissect intra-subset variability enables the identification of the specific cells that drive aging-related transcriptional signals. The model delineates core functional programs underlying each subset’s contribution to immunosenescence, providing refined biological insights into the cellular and molecular complexity of immune aging. Collectively, these findings establish IDEAL-Age as a powerful tool for dissecting immune aging at single-cell granularity, highlighting the distinct and dynamic roles of adaptive T cells, innate-like T cells, progenitors, and myeloid cells in shaping the immunological age landscape.

### Application of the IDEAL-Age model to the systemic lupus erythematosus cohort

Systemic lupus erythematosus (SLE) is a prototypical systemic autoimmune disease characterized by heterogeneous clinical manifestations and complex immunopathogenic mechanisms involving dysregulated adaptive and innate immune responses [73, 74]. Notably, accumulating evidence implicates immunosenescence, characterized by age-associated decline and remodeling of immune function, as an important contributor to SLE pathogenesis and progression [75, 76]. Given the considerable overlap between the cellular and molecular features of immunosenescence and known immunological abnormalities in SLE, leveraging a single-cell resolution aging clock to dissect immune aging dynamics in this disease offers significant promise. Precise quantification of immune aging in SLE patients could illuminate the heterogeneity of disease trajectories, identify accelerated immune aging signatures associated with flare or treatment status, and reveal novel cellular and transcriptomic biomarkers of pathophysiological relevance.

We applied IDEAL-Age to characterize aging acceleration in SLE. Following quality control, we derived the predicted ages for 156 SLE patients and 99 healthy controls from the single-cell transcriptomic profiles of their PBMC samples [76]. To evaluate the aging status of each sample, we established a benchmark using the regression curve of predicted versus chronological age in healthy samples. The residuals between the predicted and fitted ages were then employed for comparative analysis (Methods). The results indicated that managed samples exhibited a significantly accelerated aging trend, which was even more pronounced in flare samples. Notably, this trend was substantially lower in the treated group (Fig. 5a, Fig. S18).

**Fig. 5.**
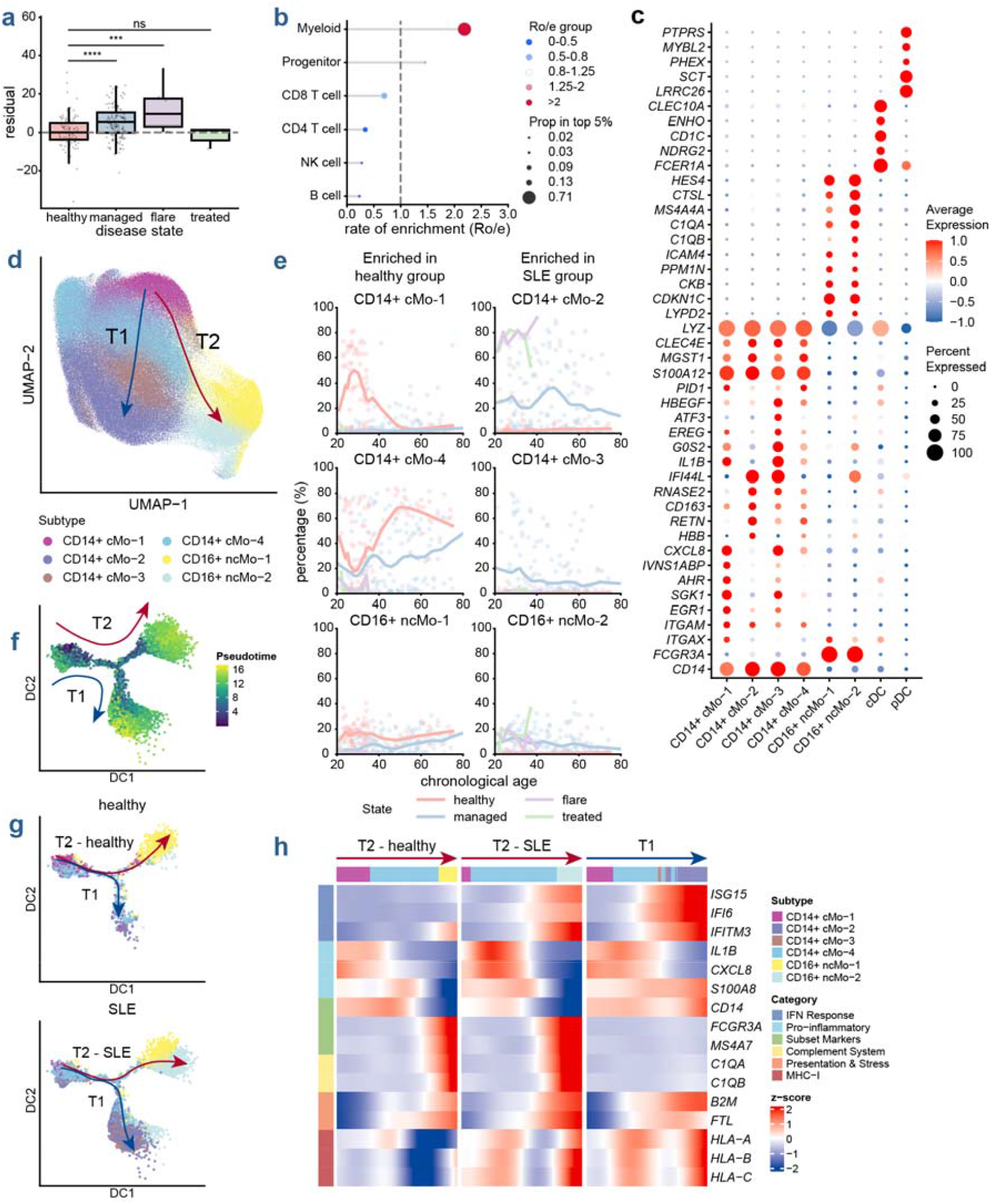
Deciphering the pathologically accelerated and divergent aging trajectories of monocytes in SLE. **a** Boxplots comparing the residuals (calculated as the difference between the predicted age and the expected age derived from the healthy cohort regression line) across healthy, managed, flare, and treated disease states. Statistical significance was determined by t-tests (ns: not significant, \*\*\**P* < 0.001, \*\*\*\**P* < 0.0001). **b** Rate of enrichment (Ro/e) of distinct cell lineages within the top 5% of highly contributing cells. The dot size represents the proportion of each lineage in the top 5% pool. The dot color indicates the Ro/e group (ranging from blue for low enrichment, ≤ 0.5, to red for high enrichment, >2). **c** Dot plot showing the average expression levels and the percentage of expressing cells for canonical marker genes across the eight identified myeloid cell subsets. **d** UMAP embedding of the monocyte compartment. Arrows indicate two distinct developmental trajectories: the T1 trajectory and the T2 trajectory. **e** Scatter plots illustrating the percentage of monocyte subsets within the total myeloid cells across chronological age in different clinical states. The bold solid lines represent the LOWESS regression curves. **f-g** Dimensional reduction projection and pseudo-time evolutionary trajectory inference of monocyte subsets based on DDRTree analysis. Arrows in **f** and **g** indicate the primary differentiation branches. Split views in g compare cell distributions between healthy controls (top) and SLE patients (bottom). **h** Heatmap displaying the dynamic gene expression patterns of genes exhibiting significant changes along the T2-healthy, T2-SLE, and T1 developmental trajectories. Cells are ordered by pseudo-time. Top color bars annotate the specific monocyte subtypes. Row annotations categorize dynamic genes into distinct functional modules.

We next examined the cell-level contributions for the healthy and SLE groups (Fig. S19a). Our analysis revealed a significant enrichment of myeloid cells among the most highly contributing cells (Fig. 5b, Fig. S19b-c). To identify the specific cellular drivers underlying this myeloid-associated aging signature, we subdivided the myeloid compartment into eight transcriptionally distinct subsets [77]: four classical monocyte subsets (CD14^+^ cMo-1 to CD14^+^ cMo-4), two non-classical monocyte subsets (CD16^+^ ncMo-1 and CD16^+^ ncMo-2), conventional dendritic cells (cDCs), and plasmacytoid dendritic cells (pDCs) (Fig. 5c-d). We found that the SLE monocyte compartment was characterized by a pronounced pathological enrichment of the CD14^+^ cMo-2 subset, accompanied by the CD16^+^ ncMo-2 subset. In contrast, healthy controls were predominantly populated by CD14^+^ cMo-1, CD14^+^ cMo-4, and CD16^+^ ncMo-1 subsets (Fig. S20a).

To understand how these disease-specific subsets evolve over the lifespan, we then tracked their proportional dynamics across chronological age (Fig. 5e, Fig. S20b). Healthy individuals exhibited a distinct age-associated transition, where the initially predominant CD14^+^ cMo-1 subset was gradually replaced by CD14^+^ cMo-4 and CD16^+^ ncMo-1 with advancing age, alongside slight expansions of the CD14^+^ cMo-2 and CD14^+^ cMo-3 populations. In contrast, SLE patients maintained persistently low levels of CD14^+^ cMo-1 across the lifespan, while demonstrating marked enrichment of the CD14^+^ cMo-2 and CD16^+^ ncMo-2 subsets. Notably, although individuals in a managed disease state displayed an age-related accumulation of CD14^+^ cMo-4 and CD16^+^ ncMo-1 that mirrored the trend in healthy controls, their absolute proportions remained significantly divergent from healthy baselines. These dynamic profiles suggest that monocytes in SLE patients follow a pathological evolutionary trajectory that is distinct from physiological aging.

To further delineate the developmental dynamics of monocytes, we performed pseudo-time trajectory analysis with Monocle2 [78] (Fig. 5d, f-g, Fig. S20c, Methods). The results revealed a bifurcation into two distinct developmental pathways originating from CD14^+^ cMo-1. The first pathway (assigned as the T1 trajectory) progresses sequentially through CD14^+^ cMo-4 and CD14^+^ cMo-3, ultimately terminating at CD14^+^ cMo-2. The second pathway (assigned as the T2 trajectory) branches from CD14^+^ cMo-4 and differentiates towards the non-classical subsets, CD16^+^ ncMo-1 or CD16^+^ ncMo-2. In healthy individuals, the T2 trajectory is predominantly favored, culminating primarily in CD16^+^ ncMo-1 (assigned as the T2-healthy trajectory). Within their T1 path, differentiation largely stalls at the CD14^+^ cMo-4 stage, with only a minor fraction reaching the terminal CD14^+^ cMo-2 state. In marked contrast, SLE patients exhibited an aberrantly hyperactivated T1 trajectory, driving a substantial proportion of cells towards the terminal CD14^+^ cMo-2 state. Meanwhile, within the T2 trajectory, monocytes from SLE patients in the managed state displayed a branching developmental fate towards either CD16^+^ ncMo-1 or CD16^+^ ncMo-2 (assigned as the T2-SLE trajectory). Notably, this terminal differentiation was heavily skewed towards a pronounced CD16^+^ ncMo-2 fate during flare and treated states.

We further dissected the dynamic gene expression patterns underlying these monocyte evolutionary trajectories (Fig. 5h, Methods). The early stages across all trajectories were characterized by high expression of classical pro-inflammatory genes (e.g., *IL1B, CXCL8, S100A8*). In the normal trajectory of healthy individuals (T2-healthy), monocytes preserved their classical characteristics and homeostatic complement functions, evidenced by high expression of *C1QA* and *C1QB*. However, as cells progressed towards the terminal states of T1 and the SLE-specific T2, there was a robust upregulation of interferon-stimulated genes (ISGs), including *ISG15, IFI6*, and *IFITM3*, along with MHC class I molecules (e.g., *HLA-A, HLA-B, HLA-C*). Specifically, cells evolving along the T1 trajectory (primarily corresponding to the abnormally enriched CD14^+^ cMo-2 subset) displayed an upregulation of pro-inflammatory cytokines and chemokines (e.g., *IL1B, CXCL8, S100A8*). Notably, while normal aging in healthy individuals involves a modest upregulation of ISGs along the evolutionary trajectory, this interferon signature is substantially upregulated at the pathological differentiation endpoints in SLE patients (Fig. S20d-e), which aligns with the well-documented “interferon signature” that is central to SLE pathogenesis [76, 79].

Taken together, these trajectory dynamics and transcriptional signatures elucidate the complex relationship between SLE and normal physiological aging. Normal aging is similarly accompanied by monocyte subset transitions and a mild accumulation of inflammatory signals. In contrast, the immune system in SLE undergoes an accelerated and pathologically divergent aging process, remodeling the transcriptional landscape of monocytes, and providing a molecular basis for systemic immune dysfunction.

Collectively, our model-derived results indicate that the characteristic premature aging phenotype in SLE arises from a pathologically divergent developmental trajectory within the myeloid compartment. Although significant ISG upregulation is also observed in T and B cells of SLE patients [80, 81], our model identifies highly inflammatory monocytes as a major contributor to their premature aging phenotype. The aberrant accumulation of hyper-inflammatory, interferon-responsive terminal monocyte subsets alters the immune landscape, thereby contributing to this accelerated immune aging and systemic dysfunction.

## Discussion

In this study, we present IDEAL-Age, an immune aging prediction framework that integrates multi-cohort single-cell transcriptomic data from four independent PBMC datasets. Our model delivers not only robust immunological age prediction but also multi-scale interpretability spanning gene- and single-cell-level contributions for each donor, thereby advancing the resolution at which immunosenescence can be characterized. This interpretability addresses the limitations of bulk or pseudo-bulk approaches by capturing intra-population heterogeneity and revealing discrete transcriptomic signatures underlying immune aging at single-cell granularity.

A significant finding of this study is the identification of non-monotonic aging signatures, such as U-shaped and inverted U-shaped gene contribution trajectories. Traditional molecular clocks, rooted in linear regression, inherently prioritize genes with constant rates of change, often overlooking critical phase-specific biological transitions. Our model demonstrates that biological aging is not a purely linear decline but a series of coordinated shifts. By aligning computationally derived inflection points with established proteomic waves and endocrine milestones (e.g., the menopausal transition), we show that deep learning can capture the ‘rise and fall’ of biological priorities across the lifespan. This resolution reveals a layered architecture of immunosenescence, where adaptive immune signals are gradually superseded by innate-dominant inflammatory markers and systemic proteostatic stress.

The observed numerical disparity between U-shape and inverted U-shape genes suggests that our approach delineates underlying biological trajectories rather than methodological artifacts. The larger proportion of U-shape genes may reflect the systemic dysregulation of homeostasis and subsequent compensatory stress responses during aging, whereas the more constrained set of inverted U-shape genes likely signifies age-specific regulatory milestones associated with mid-life physiological transitions. Notably, the identified peak ages within the inverted U-shape sub-clusters closely align with the three major non-linear shifts in systemic aging previously reported in plasma proteomics [58], implying potential cross-omic synchrony during key chronological transitions over the human lifespan.

Importantly, the single-cell resolution offered by IDEAL-Age improves upon the granularity of prior pseudo-bulk models by enabling the dissection of cell-subset-specific as well as intra-subset heterogeneity in aging contributions. Our analyses revealed consistent and statistically significant enrichment of age-predictive signals within specific subsets such as CD8^+^ naïve T cells, MAIT cells, and hematopoietic progenitors, with functional enrichment in pathways reflecting both immune surveillance and cellular biosynthesis. The heterogeneity in gene- and cell-level contributions across datasets further highlights the complex influence of cohort-specific biological and technical factors, reinforcing the necessity for adaptable and interpretable modeling frameworks to capture diverse aging patterns.

In this study, we found that the elevated immune age in SLE is largely associated with inflammation-associated monocytes. These cells, characterized by aberrantly amplified pro-inflammatory and interferon responses, not only serve as the core predictive features for the model but also substantially deplete the systemic adaptive immune reserve. Meanwhile, we found that their pathogenic trajectory exhibited a negative correlation with CD4^+^ and CD8^+^ naïve T cell abundance, suggesting that continuous inflammatory stimulation may contribute to the eventual exhaustion of reserve-maintaining T cells. Furthermore, the predicted immune age notably decreased in clinically treated patients, which closely paralleled a significant recovery of CD8^+^ naïve T cells. This provides a possible explanation for the reduction in the predicted immune age following treatment.

Despite the robust performance of IDEAL-Age, several limitations warrant consideration. First, while we successfully implemented a targeted fine-tuning strategy to mitigate the initial predictive biases at demographic extremes (<19 and >77 years), the current scarcity of large-scale, strictly independent out-of-distribution scRNA-seq cohorts for pediatric and supercentenarian populations restricts our capacity for extensive cross-cohort validation in these specific age groups. Consequently, fully establishing the model’s robust generalizability for early developmental or extreme longevity studies remains an ongoing objective. Additionally, batch effects and technical variability intrinsic to scRNA-seq technologies pose challenges in harmonizing data across cohorts, potentially influencing model generalizability. Certain rare or transcriptionally highly variable immune cell types may be underrepresented, impacting the granularity and completeness of cell-level contributions. Finally, transcription-based aging assessments, while informative, represent only one dimension of immune senescence; integration with epigenomic, proteomic, and functional immune phenotypes remains essential for a comprehensive characterization.

Future efforts should focus on integrating emerging large-scale, age-extreme cohorts to further validate and refine the model’s boundary predictions across broader demographic landscapes. Longitudinal sampling and integration of multimodal single-cell techniques will enhance biological insights into cellular aging and intercellular crosstalk. Extending validation to ethnically and environmentally diverse populations, as well as across other age-related diseases, will further facilitate clinical translation. Additionally, as high-quality, longitudinal single-cell datasets from human clinical anti-aging intervention trials become available, deploying our method to track molecular rejuvenation signatures will be a key direction for future research. Through such comprehensive approaches, IDEAL-Age can evolve into a broadly applicable computational tool for precision immunogerontology, guiding targeted interventions to mitigate immunosenescence and promote healthy aging.

## Conclusion

We present IDEAL-Age, a single-cell transcriptome-based immune aging model with multi-scale interpretability that advances our understanding of immunosenescence. The model robustly predicts immunological age across diverse cohorts, revealing both conserved and dataset-specific gene signatures and non-linear transcriptomic dynamics that align with physiological aging waves. Single-cell resolution uncovers heterogeneous aging states even within cell subsets, particularly highlighting hematopoietic progenitors and T cells. Application to systemic lupus erythematosus suggests accelerated immunological aging associated with interferon-associated monocyte shifts, underscoring clinical relevance and utility for studying immunosenescence.

## Methods

### Data sources

Seven publicly available single-cell RNA-sequencing datasets of human peripheral blood mononuclear cells (PBMCs) were utilized in this study: the AIDA, OneK1K, siAge, SC2018, HCA, Sound Life, and CIMA cohorts. Donor-level metadata, including chronological age, sex, and clinical annotations, were obtained from the original files or publications and standardized to a unified format to ensure consistency across cohorts.

### Feature selection for a robust and biologically informative gene panel

Prior to model training, we implemented a comprehensive feature selection pipeline to identify a robust and biologically meaningful set of genes, thereby reducing data dimensionality and mitigating model overfitting. This pipeline integrated three independent criteria to ensure the selected genes were informative for immunological age prediction.

First, to capture fundamental cellular heterogeneity, we identified cell-type marker genes by performing differential expression analysis across major immune cell subsets. To ensure a comprehensive and representative feature space, we performed unified cell-type annotation across our training cohorts using SCimilarity. This process resulted in the identification of 11 major cell types: CD4^+^ T cell, CD8^+^ T cell, NK cell, monocyte, B cell, macrophage, platelet, DC, plasma cell, ILC, and mast cell. Based on these unified annotations, we employed our previously developed algorithm, eMark [82], to identify marker genes for each of the 11 cell types. Specifically, we selected genes with a weight greater than 0.1 as cell-type-specific marker genes. This procedure yielded a final refined feature space consisting of 2,425 unique genes. This gene set was utilized as part of the input features for our aging clock model, ensuring that the model focuses on biologically relevant variation across the major immune compartments.

Second, to directly select features with strong statistical association with age, we performed univariate feature selection based on mutual information (MI). We computed the MI between each gene’s expression level and donor chronological age across all our training cohorts. The top 2,000 genes with the highest MI scores were selected as highly age-informative.

Third, we incorporated eight publicly available aging-related gene sets (SenMayo, CellAge, GenAge, ASIG, SASP, AgingAtlas, SenUp, and SigRS) from curated databases including MSigDB and the Human Ageing Genomic Resources (HAGR). This allowed the inclusion of genes with prior evidence linking them to biological processes of aging.

Finally, we took the union of the gene sets derived from the above three criteria to form a comprehensive feature list (Fig. S1). We then intersected this gene set with the features present in our single-cell datasets and filtered out 75 genes that were not detected, resulting in a final feature space of 5,012 genes for all subsequent model training and validation. At the individual-cell level, genes with no detected expression were naturally retained as zeros within the sparse raw count matrix. To ensure numerical robustness during model training, any abnormal non-finite values (e.g., NaN or inf) were explicitly zero-filled during the data loading pipeline prior to downstream z-score normalization.

### Cohort definition and partitioning

The complete cohort was partitioned into training, internal test, and external test sets to ensure rigorous assessment of generalization capability. The training cohort comprised combined AIDA and OneK1K training splits (1,284 donors) and was used exclusively for model parameter optimization. Internal test cohorts consisted of the AIDA test split (n=125) and OneK1K test split (n=197), both withheld during training to provide unbiased evaluation on data from the same generation protocols. To enhance the generalizability of our model across the human lifespan, we incorporated additional diverse cohorts for model fine-tuning, including siAge (n=61), comprising neonates and children, and SC2018 (n=7), featuring centenarians. External test cohorts included HCA (n=48), Sound Life (n_donor_=96, n_sample_=255) and CIMA (n=421), which served as independent datasets for assessing cross-study generalization capability under different laboratory conditions, sequencing protocols, and demographic distributions.

Donor availability was verified through automated file system traversal, identifying all donors with complete H5AD files and non-missing chronological age metadata. The final cohort spanned ages 0–110s years, with age distribution characteristics (mean ± standard deviation) as follows: AIDA training (41.4 ± 12.4 years), AIDA test (40.5 ± 12.5 years), OneK1K training (64.0 ± 16.6 years), OneK1K test (63.7 ± 16.1 years), siAge (32.6 ± 29.3 years), SC2018 (110s years), HCA (43.3 ± 14.9 years), Sound Life (45.7 ± 14.6 years) and CIMA (35.3 ± 13.3 years). Specifically, AIDA contained only 2 donors under age 20 and no donors over age 80, while OneK1K contained 5 donors under age 20 and 129 donors over age 80.

### Data preprocessing

#### Pseudo-bulk aggregation

For baseline models operating on fixed-size feature vectors, donor-level pseudo-bulk expression profiles were constructed by averaging gene expression across all cells within each donor. For each donor *d* with *N*_*d*_ cells, the pseudobulk expression vector 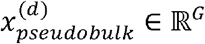 was computed as:

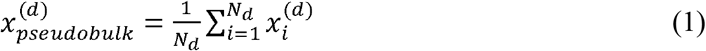

where 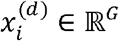 denotes the expression vector of cell *i*. Raw counts from layers[‘counts’] were prioritized when available to preserve count-based statistical properties; otherwise, log-normalized values from X were utilized. This aggregation was performed independently for each donor to prevent information leakage between training and test sets.

#### Single-cell data preparation

For deep learning architectures designed to leverage cellular heterogeneity, raw single-cell expression matrices were retained without aggregation. To balance computational feasibility with information preservation, preprocessing steps were applied to manage the variable-length nature of single-cell datasets while maintaining biological fidelity.

Donors with *N*_*d*_ > *M* cells underwent random subsampling without replacement to retain exactly *M* cells, where *M* represents the maximum cells per donor hyperparameter. The subsampling procedure was implemented as follows: if the donor’s cell count *N*_*d*_ was less than or equal to *M*, the original expression matrix was retained without modification; otherwise, a uniform random sample of size *M* was drawn from the available cell indices, and the corresponding rows were extracted to form the subsampled matrix. Multiple values of *M* spanning {100, 500, 1000, 10000} were evaluated to characterize the accuracy-efficiency trade-off, with detailed analyses presented in Supplementary Methods. Based on this evaluation, *M =*1000 was selected as the default configuration for subsequent analyses, providing an optimal balance between computational tractability and information content. Importantly, while the pseudo-bulk baseline features were constructed by aggregating all available cells (*N*_*d*_) from a donor, the DeepSets-Attention architecture strictly utilizes this restricted subsample of *M* cells. This means our IDEAL-Age model achieves its superior predictive performance despite utilizing a limited subset of cells, highlighting the right informational value of single-cell resolution compared to the average signals of the entire cell population.

To stabilize neural network optimization and mitigate batch effects across datasets, z-score normalization was applied at the gene level. For each gene *g*, the mean *μ*_*g*_ and standard deviation *σ*_*g*_ were estimated from a randomly selected subset of 50 training donors:

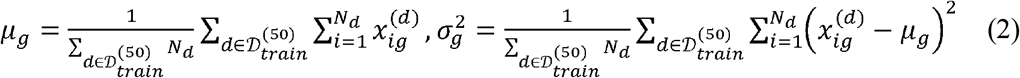

where 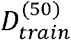 denotes the random subset. This subset size was chosen to balance statistical robustness (approximately 50,000 cells total when *M =*1000) with computational efficiency, ensuring stable parameter estimates without requiring full-dataset traversal. The transformation was then applied uniformly to all cells:

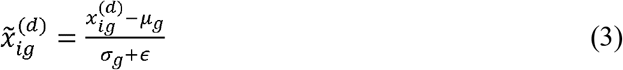

with *ϵ* = 10^−6^ included to prevent division by zero for genes exhibiting invariant expression patterns.

#### Model architecture

Existing transcriptomic aging clocks based on bulk RNA-seq or pseudobulk aggregation suffer from a fundamental limitation: averaging expression across millions of cells masks the considerable immunophenotypic heterogeneity that fundamentally defines immunosenescence. Single-cell studies have revealed that aging-associated changes exhibit marked cell-type specificity. For instance, CD8^+^ CD28^-^ senescent T cells progressively accumulate with age, whereas naïve CD4^+^ T cells decline. These opposing trajectories within T cell subsets cannot be captured by bulk measurements, which conflate signals across all cell types. To preserve this granular information and enable mechanistic interpretation, we developed a hierarchical deep learning framework that operates directly on single-cell expression profiles.

The architecture design was guided by three fundamental principles derived from the mathematical properties of single-cell data. First, the model must exhibit permutation invariance, ensuring that predictions remain unchanged regardless of the arbitrary ordering of cells within scRNA-seq datasets. Second, the architecture must accommodate variable set sizes, as donor cell counts typically range from 500 to 10,000 cells post-quality control. Third, the model should provide biological interpretability by exposing cell-level and gene-level contributions amenable to downstream biological validation.

#### DeepSets-Attention architecture

The complete architecture, termed DeepSets-Attention, comprises three sequential modules that hierarchically process information from the gene expression level through cellular representations to donor-level age predictions.

The first module implements a cell-level encoder *ϕ*: ℝ^*G*^ ⟶ ℝ^*d*^ that independently maps each cell’s expression profile to a *d*-dimensional latent representation. The encoder *ϕ* was implemented as a 3-layer multilayer perceptron (MLP) with interleaved normalization and nonlinearity:

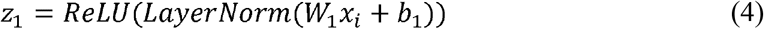

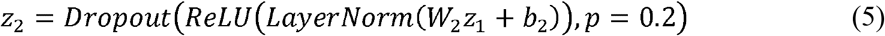

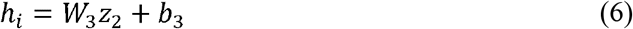

Layer dimensions progress as *G* ⟶ *h*_*hidden*_ ⟶ *h*_*hidden*_ ⟶ *d*, where *h*_*hidden*_ = 1024 and *d =*256. LayerNorm was applied before nonlinearities to stabilize gradients during training. Critically, encoder parameters *θ*_*ϕ*_ = {*W*_1_,*b*_1_,*W*_2_,*b*_2_,*W*_3_,*b*_3_} are shared across all cells, enforcing the permutation equivariance property: applying the encoder to a permuted cell set yields a correspondingly permuted encoding set.

The second module implements multi-head attention pooling to aggregate cellular information while capturing the multifaceted nature of immunosenescence. Traditional neural network architectures employ simple summation or mean pooling to achieve permutation invariance. However, immunosenescence manifests through multiple independent processes—including thymic involution, T cell exhaustion, and inflammaging—each involving distinct cell subsets. To capture these diverse biological signatures, we employ a multi-head attention pooling mechanism inspired by set-based representation learning. The pooling module learns *r* latent query vectors *q*_1_, …, *q*_*r*_ ∈ ℝ^*d*^ that adaptively attend to relevant cells.

The attention operation is formally defined as:

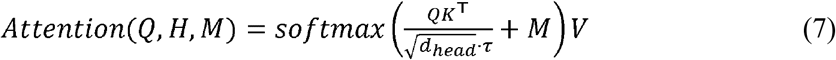

where *Q = Q*_*latent*_*W*_*Q*_ is the query projection with *W*_*Q*_ ∈ ℝ^*d*×*d*^ : *Q*_*latent*_ ∈ ℝ^*r*×*d*^ represents the learnable initial query matrix initialized from 𝒩 (0,0.02^2^); *H* = [*h*_1,_ …, *h*_*N*_]^T^ ∈ ℝ^*d* ×*d*^ denotes the cell encoding matrix; *K* = *HW*_*K*_ and *V* = *HW*_*V*_ are key and value projections with *W*_*K*,_ *W*_*V*_ ∈ ℝ^*d*×*d*^ *d*_*head*_ =*d*/*h* represents the per-head dimensionality with *h* =4 attention heads; *τ* >0 is a temperature parameter controlling attention sharpness (default *τ* =1.0); and *M* ∈ {0, −∞}^1×*N*^ is a binary mask assigning *M*_*i*_ = − ∞ to padded (invalid) cells.

To enable multi-head attention, projections are reshaped into head-specific subspaces. Specifically, the projected queries *Q* are partitioned into multi-headed projected queries as 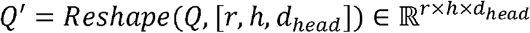 with analogous transformations for keys and values. Attention logits are computed per-head as 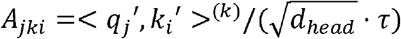, where <·,·>^(*k*)^ denotes the dot product for head *k*. After applying the mask and softmax normalization to obtain attention weights *α*_*jki*_, the pooled representations are computed as 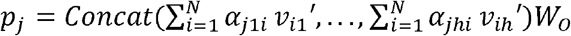, where *W*_*o*_ ∈ ℝ^*d*×*d*^ is an output projection matrix.

Each query vector learns to specialize in detecting specific cell populations or functional states through end-to-end training. This specialization emerges without explicit supervision, driven solely by the age prediction objective.

The third module implements a donor-level prediction head *ρ*: ℝ^*d*^ ⟶ ℝ that maps aggregated cellular representations to age predictions. The *r* query-specific representations are first combined through mean pooling, then processed through a 3-layer MLP with progressive dimensionality reduction:

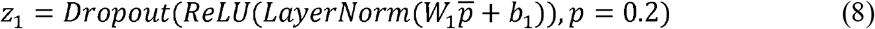

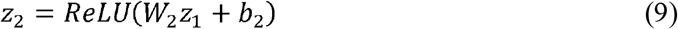

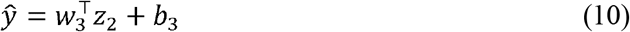

where 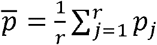 and layer dimensions progress as *d* ⟶ 512 ⟶ 256 ⟶ 1.

The complete architecture satisfies permutation invariance: predictions remain unchanged when cells are reordered, as cell ordering does not affect the encoded set {*h*_*i*_}, attention is computed over the unordered set with softmax normalization, mean pooling over queries is order-independent, and the final MLP operates on a single pooled vector.

### Multi-task regularization

To improve age prediction accuracy while providing interpretable age range estimates, an auxiliary classification task was introduced alongside the primary regression objective. Chronological ages were discretized into *K* =10 uniform bins spanning 10-year intervals: *B*_*k*_ = [10*k*, 10(*k* + 1)) years for *k* = 0,…,9. A separate classification head predicts the probability distribution over these age bins via Softmax transformation of the pooled representation. The composite loss function combines three objectives:

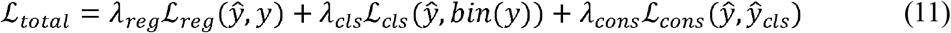

The regression loss employs Smooth L1 (Huber loss), which transitions from quadratic to linear behavior for large errors, providing robustness to outliers. The classification loss uses standard cross-entropy on the age bin predictions. The consistency loss ℒ_*cons*_ enforces agreement between the direct continuous age regression prediction 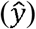 and the expected age implied by the categorical classification distribution 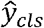. It is formulated as:

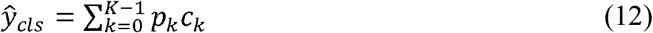

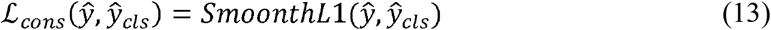

where *p*_*k*_ is the predicted probability for the *k*-th age bin *B*_*k*_, *c*_*k*_ = 10*k* + 5 is the centroid of that bin. This formulation ensures that the model’s categorical understanding of age groups is strictly consistent with its precise continuous predictions.

### Ensemble training strategy

#### AutoGluon framework integration

Rather than training DeepSets-Attention in isolation, we integrated it into an ensemble of diverse predictors using AutoGluon-Tabular, a framework that automates model selection, hyperparameter optimization, and ensemble construction. This approach offers three key advantages over single-model training. First, AutoGluon evaluates multiple baseline model families—including feedforward networks, neural networks, and k-nearest neighbors—automatically selecting the best-performing subset for each dataset. Second, the framework employs stacked ensembling, where out-of-fold predictions from base models serve as meta-features for a higher-level model, often surpassing simple weighted averaging. Third, Bayesian optimization automatically tunes hyperparameters across the model space, eliminating manual tuning while providing principled exploration of the configuration landscape.

#### Two-stage training protocol

Model training proceeded through two sequential stages designed to first establish a strong baseline ensemble, then augment it with custom single-cell models. In Stage 1, AutoGluon’s TabularPredictor was invoked with pseudobulk expression features to train baseline models including FastAI tabular learner (a feedforward network with automatic architecture search) and PyTorch feedforward networks (standard MLPs with configurable depth). These models were trained using 5-fold cross-validation within a 10-minute time budget, with validation mean absolute error guiding model selection and retention.

In Stage 2, DeepSets-Attention model was registered as an AutoGluon-compatible predictor by subclassing AbstractModel and implementing the required interface methods for training, prediction, default hyperparameters, and resource requirements. Single-cell data were shared across cross-validation folds via class-level attributes, allowing the custom models to access raw cellular information while maintaining compatibility with AutoGluon’s tabular interface. When added to the ensemble via fit_extra(), AutoGluon automatically aligned cross-validation splits with Stage 1 folds, generated out-of-fold predictions for stacking, and optimized ensemble weights through greedy forward selection or linear regression on the validation predictions.

The complete training procedure for DeepSets-Attention models within each fold is detailed in Algorithm 1. After initializing model parameters and fitting a StandardScaler on a random subset of 50 training donors, training proceeds through multiple epochs. In each epoch, the training donors are shuffled and partitioned into batches. For each batch, individual donors are loaded, normalized, potentially subsampled if exceeding *M* cells, padded to uniform length, and stacked into batch tensors. These batches are then processed through the cell encoder, attention pooling, and prediction heads to compute age predictions and auxiliary classification outputs. Losses are calculated, backpropagated, and parameters are updated via gradient descent. This procedure repeats until convergence, yielding trained models suitable for ensemble integration.

#### Hyperparameter configuration

Hyperparameters for the DeepSets-Attention model were determined through a combination of prior work, preliminary experiments, and AutoGluon’s automated hyperparameter optimization. The cell encoder employs a hidden dimension of 1024 to provide sufficient capacity for the approximately 5,000 genes, compressing the representation to an output dimension of 256 for efficient downstream processing. Dropout regularization with a probability of 0.2 was applied uniformly across encoder and prediction head layers. The attention pooling module uses 4 learnable query vectors to capture major immune aging axes, with 4 attention heads providing multi-perspective aggregation and a temperature parameter of 1.0 balancing attention sharpness. The prediction head progressively compresses information through hidden dimensions of 512 and 256 before final age prediction. For the auxiliary multi-task objective, 10 age bins spanning 10-year intervals were used with loss weights of 0.5 and 0.1 to moderately regularize the primary regression task. Optimization employed the Adam algorithm with a learning rate of 0.001, a batch size of 8, and 105 epochs allowing empirical convergence. The cell subsampling threshold was set to 1000 cells per donor based on accuracy-efficiency trade-off analysis.

AutoGluon’s Bayesian hyperparameter search explored key architectural choices including hidden dimensions in powers of 2 from 16 to 1024, learning rates sampled log-uniformly from 0.0001 to 0.1, and dropout probabilities uniformly sampled from 0.0 to 0.5. A total of 20 trials were conducted with 5-fold cross-validation per trial, selecting the configuration that minimized validation MAE.

##### Algorithm 1 DeepSets Training Procedure

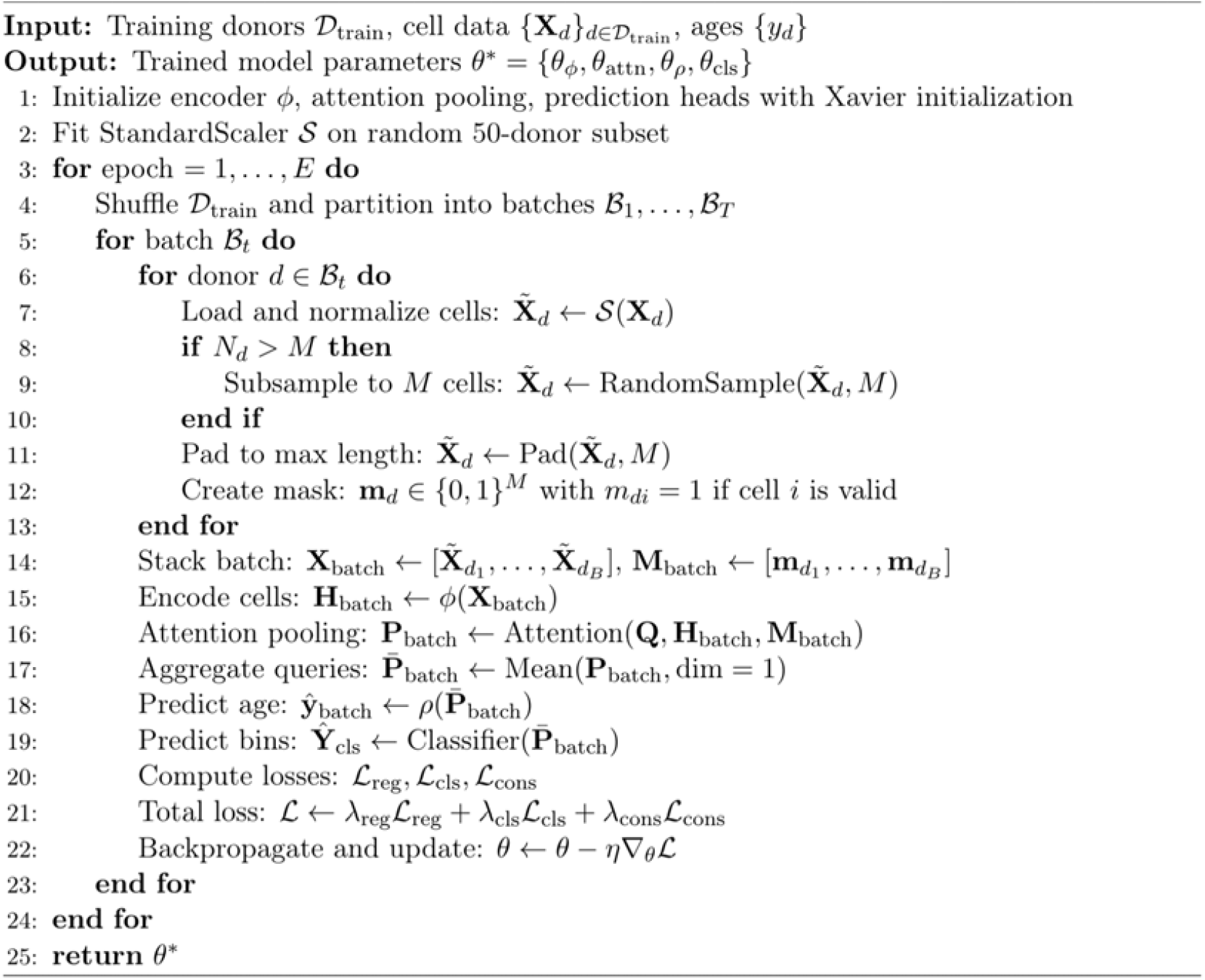

### Model interpretability methods

#### Motivation for multi-method approach

Interpretability is essential for clinical translation and biological discovery in aging research. However, different attribution methods capture complementary aspects of model behavior: gradient-based methods quantify sensitivity to perturbations, attention weights reveal learned importance patterns, and integrated gradients provide axiomatic guarantees of attribution completeness. Rather than relying on a single technique, we implemented six complementary methods to triangulate cell-level and gene-level contributions from multiple theoretical perspectives.

#### Cell-level attribution methods

All attribution methods assign a scalar contribution *s*_*i*_ ∈ ℝ to each cell *i* within a donor. To facilitate cross-donor comparison and visualization, contributions are normalized to the [0, 1] interval per donor. The six methods span gradient-free and gradient-based approaches with varying computational costs and theoretical properties.

While our main narrative primarily highlights the attention-based attribution for cell-level interpretation—due to its intuitive capacity to capture how the model weights specific biologically relevant cell subsets—the additional five methods are fully integrated into the software package. We provide this diverse suite of attribution methods as a versatile interpretability toolkit for the community, empowering users to easily conduct comparative analyses and select the most appropriate interpretability perspective (e.g., gradient-based sensitivity versus activation-based feature magnitude) for their specific datasets and research contexts. The attention mechanism naturally provides contributions without additional computation. For a donor with *N* cells, the attention matrix *A* ∈ ℝ^*r×h×N*^ contains weights *α*_*jki*_ representing the attention from query *j* and head *k* to cell *i*. Cell importance is quantified by averaging across all queries and heads: 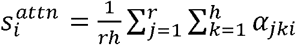. Cells with high attention contributions are consistently prioritized across multiple queries, suggesting broad biological relevance across different aging dimensions.

Activation magnitude provides an alternative gradient-free measure by computing the L2 norm of each cell’s encoded representation: 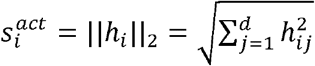. This contribution identifies cells with distinct expression profiles that produce large encoder activations, potentially corresponding to rare or extreme cell states.

Gradient-based sensitivity analysis quantifies how prediction changes with respect to cell encodings through a first-order Taylor expansion: 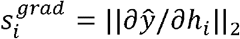. Cells with high gradient magnitudes exert a strong influence on predictions, such that small perturbations to their representations significantly alter the predicted age. Gradients are computed via backpropagation with the predicted age as the scalar loss.

The gradient-input method combines sensitivity with feature magnitude through elementwise multiplication 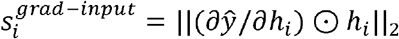. This approach balances “how sensitive” (captured by gradients) with “how large” (captured by activations), proving particularly useful when encoder outputs exhibit wide dynamic range across cell types.

Integrated gradients (IG) provide a principled attribution satisfying axiomatic properties including sensitivity and completeness. For computational efficiency, we compute IG in the encoded space *H* rather than input space *X*. The method integrates gradients along a linear path from a baseline 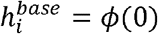 (encoding of zero expression) to the actual encoding *h*_*i*_. The integral 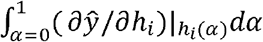 is approximated via Riemann sum with 32 steps, where 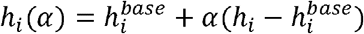 defines the interpolation path. The resulting attribution 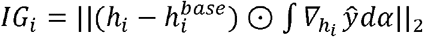 satisfies the completeness axiom: attributions sum to the prediction difference from the baseline.

For maximum granularity, IG can be computed directly on gene expression in the input space, yielding cell-by-gene attribution matrices *IG*_*ig*_ that quantify each gene’s contribution in each cell. Cell-level contributions are then obtained by summing across genes: 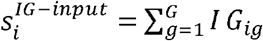. While this provides the finest-grained attribution, it requires *K × B* forward and backward passes (where *K* = 32 is the number of integration steps and *B* is the batch size), making it computationally expensive and best reserved for detailed case studies.

### Gene-level attribution methods

Gene importance contributions *c*_*g*_ ∈ ℝ quantify each gene’s aggregate contribution to age prediction across all cells within a donor. Three complementary methods were implemented mirroring the cell-level approaches. Pure gradient attribution computes 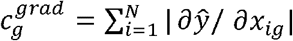, aggregating the magnitude of prediction sensitivity to each gene across all cells. The gradient-input variant 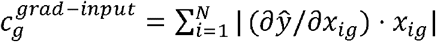 downweights genes with low expression despite high gradients, focusing attribution on genes that are both sensitive and actively expressed. Integrated gradients 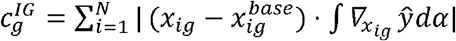 provides the most principled gene attribution through path integration from zero baseline expression, satisfying completeness and ensuring attributions sum to the prediction difference.

### Evaluation framework

#### Performance metrics

Model performance was assessed using four complementary metrics capturing different aspects of prediction quality. The Pearson correlation coefficient (PCC) measures linear association between predicted and chronological ages through the formula 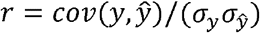, providing a scale-free measure of correlation strength. Mean absolute error (MAE) quantifies the average prediction error in years as 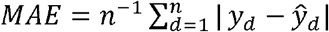, providing an intuitive measure of typical deviation.

#### Cross-dataset evaluation protocol

To rigorously assess generalization capability, all models were evaluated on both internal and external validation sets without any retraining or fine-tuning on the test data. The internal test sets comprising the AIDA and OneK1K test splits share data generation protocols with their corresponding training splits, providing an evaluation under matched experimental conditions. The external validation sets consisting of the HCA, Sound Life, and CIMA datasets originate from different laboratories with distinct sequencing protocols and demographic compositions, offering an assessment of cross-study generalization under realistic deployment conditions. All hyperparameters were fixed prior to evaluation based solely on training and internal validation performance.

#### Benchmark methodology for aging markers, gene sets, and scoring metrics

Pseudo-bulk data for individual aging markers and gene sets were generated using the scanpy.get.aggregate function from the Scanpy Python package, with the aggregation parameter set to by=‘sum’ and layers=‘counts’. For the aging marker and gene set methods, expression values were then normalized and log-transformed using the scanpy.pp.normalize_total and scanpy.pp.log1p functions. For individual senescent gene markers, we calculated the absolute values of the PCC between the pseudo-bulk expression levels and chronological age across all samples. For senescence-associated gene sets, the average expression levels were quantified using the scanpy.tl.score_genes function. The resulting scores were then correlated with age.

#### PLSR (cell composition) implementation

We employed SCimilarity for the unified annotation of cell subsets. The specific cell types and their corresponding abbreviations are detailed in Table S8. The proportions of all cell subsets within each sample were utilized as features for training the aging clock model. The partition of training and testing sets remained consistent with the distribution used for our IDEAL-Age model. Following the original implementations [24], the PLSR model was constructed using sklearn.cross_decomposition.PLSRegression with n_components=3.

#### scImmuAging implementation

For the scImmuAging benchmarking, SCimilarity was used for the unified annotation of major cell lineages (see Table S8). In accordance with the original study’s methodology [25], we specifically selected B cells, NK cells, CD4^+^ T cells, CD8^+^ T cells, and monocytes to evaluate the performance of their respective cell-type-specific models.

#### Ablation test

Ablation studies were conducted to evaluate model performance across multiple dimensions. For the benchmarking of single-cell, bulk-level, and ensemble models, we directly compared their respective predictive outputs. To assess the impact of feature selection, we retrained a standalone model based on the global gene set (full feature space) for comparison. Given that the training sets were all sequenced using the 10x Genomics single-cell platform, we utilized the official 10x Genomics GRCh38 2024-A reference genome. We defined the ‘global gene set’ as the union of all genes mapped within the AIDA and OneK1K datasets; any missing values relative to this global set were imputed with zeros. For all the test sets, only genes mapping to this global set were retained, with missing genes similarly zero-filled to serve as input for the global-gene model.

### Downstream analysis

#### Dataset usage

For the downstream analysis, we utilized four datasets including AIDA, OneK1K, HCA, and siAge. Given the large scale of the Sound Life and CIMA datasets, they were used solely as external benchmarks rather than for downstream applications, primarily due to computational constraints and memory loading limitations in R.

#### Data alignment

To ensure feature consistency across all datasets, we defined a reference set consisting of 19,808 protein-coding genes identified in the training cohort. For all other datasets, only these 19,808 genes were retained. Any genes missing from a specific dataset were zero-filled to maintain a uniform input dimensionality.

#### Gene contribution evaluation across datasets

We utilized integrated gradients (IG) to quantify gene importance in our analysis. IG was selected as the definitive metric due to its superior mathematical guarantees for biological interpretability: it satisfies both sensitivity (evaluating the attribution path from a zero-expression biological baseline rather than relying solely on local gradients) and completeness (ensuring the sum of attributions accurately equals the difference between the model’s predicted age and the baseline prediction).

#### Functional enrichment analysis

Functional enrichment analysis was performed using the enrichr function from the Python package GSEAPY, based on the provided gene lists. The parameters were configured with organism=“Human” and cutoff=0.05. The gene sets utilized for the functional enrichment analysis included GO_Biological_Process_2025, KEGG_2021_Human, and MSigDB_Hallmark_2020. For the analysis of the U-shape genes and inverted U-shape genes, Reactome_Pathways_2024 was also utilized.

For the analysis of the top 10% contributing genes, the top five most significant functional terms within each dataset (ranked by *P*-value significance) were extracted for each evaluated gene set. These selected terms were then aggregated to generate a non-redundant union set of highly enriched pathways across all datasets. To construct a comparative matrix, the corresponding *P*-values for these union terms were retrieved from each dataset. For terms absent from a specific dataset’s results, a non-significant default *P*-value of 1 was imputed. All *P*-values were subsequently -log_10_ transformed. Finally, terms were grouped by their overarching gene sets and ranked based on their maximum -log_10_(*P*-value) across the four datasets.

#### Trend pattern analysis of gene contribution across age

To characterize the developmental trajectories of gene contributions across the aging process, we implemented a trend classification algorithm that first applies LOWESS to capture underlying patterns while reducing noise, then calculates first-order differences of the smoothed values to assess local trend directions. Using a tolerance threshold of 10% to accommodate biological variability, we classified trends as monotonically increasing if ≤10% of differences were negative, monotonic decreasing if ≤10% were positive, and identified single-inflection patterns (up-then-down or down-then-up) based on sign changes in the derivatives, with complex multi-inflection patterns categorized as “others”, thereby systematically capturing five distinct trajectory types in gene contribution dynamics across the lifespan (Fig. 3c, Fig. S10).

#### Determination the precise turning points for non-linear age-related trends

LOWESS was applied to determine the precise turning points (the “inflection ages”) for genes characterized by non-linear age-related trends (U-shape and inverted U-shape). For each gene, the mean contribution across chronological age was modeled using the loess function in R with a span parameter of 0.75. To accurately identify the inflection point, we generated a dense grid of 500 equally spaced age points spanning the minimum to maximum chronological age of the dataset and predicted the corresponding contribution values using the fitted LOWESS model. The inflection age was defined as the age at the minimum predicted value (trough) for U-shape genes, and the age at the maximum predicted value (peak) for inverted U-shape genes. Genes with fewer than five valid data points were excluded from this calculation.

#### Definition of the subgroups of the U-shape and inverted U-shape groups

To identify distinct patterns within each non-linear age-related trend (U-shape and inverted U-shape), we performed unsupervised 1-dimensional *k*-means clustering on the calculated inflection ages. Clustering was performed separately for each trend group using the k-means algorithm with 25 random initial configurations (nstart = 25). Based on the underlying data distributions, we specified *k* = 3 for inverted U-shape genes and *k* = 4 for U-shape genes.

To visualize the aggregated dynamics of each subgroup, individual gene trajectories were overlaid with a group-level smoothed curve, calculated via LOWESS regression (span = 0.75) on the pooled data of all genes within the cluster. Group-specific peak or trough coordinates were determined by predicting values across a simulated age grid (20 to 100 years, length = 500) using the group-level LOWESS model. All data visualization, including density plots and faceted trajectory graphs, was conducted in R using the ggplot2 and patchwork packages.

### Statistical modeling of sex-specific trajectories

To robustly test for sex-based divergences in nonlinear aging trajectories, Generalized Additive Models (GAMs) were constructed using the mgcv package in R. The mean contribution was modeled as a function of chronological age using thin plate regression splines (bs = “tp”). To test for overall sex differences, a likelihood ratio test (Chi-square) was performed comparing a null model (assuming a single consensus aging trajectory for both sexes) against an interaction model (incorporating sex as a parametric term and sex-specific smooths: s(true_age, by = sex)). The resulting *P*-values were adjusted for multiple comparisons across all sub-groups using the Benjamini-Hochberg (FDR) procedure.

### Estimation of turning points

To explicitly identify the critical “turning points” (the age of trajectory reversal), we evaluated the first derivatives of the fitted GAM smooths utilizing the gratia package in R. True turning points for the U-shape and Inverted U-shape trajectories were defined as the chronological age at which the first derivative of the sex-specific smooth crossed the zero axis (transitioning from negative to positive, or vice versa).

### Uniform cell type annotations

To ensure uniform cell type annotation, we used the MapQuery function from the R Seurat package to map the cell type annotations of AIDA, HCA, and siAge based on the predefined annotations of OneK1K dataset (“predicted.celltype.l2”).

### Definition of accelerated and decelerated aging

We defined accelerated and decelerated aging based on the residuals of predicted age within each cohort. Specifically, we applied LOWESS to the healthy samples within each cohort to establish a regression curve between chronological and predicted age. For each individual, the age residual was calculated as the difference between the predicted age and its corresponding value on the fitted regression curve. Within each cohort, we determined the standard deviation of these residuals across all samples. Accelerated, normal, and decelerated aging were subsequently defined by the relationship between an individual’s residual and plus/minus one standard deviation.

### Identification of putative pro-aging and pro-youthful cellular phenotypes

To identify cell subsets associated with accelerated or decelerated aging across the AIDA and OneK1K datasets, we evaluated both the shifts in their relative proportions (log_2_FC) and their predictive contribution in IDEAL-Age models. A cell subset was retained for further analysis only if its model contribution differed significantly from the normal baseline (*P* < 0.05). To ensure biological relevance, we strictly excluded discordant trends, specifically filtering out subsets that exhibited unidirectional proportional shifts (i.e., simultaneous expansion or depletion in both accelerated and decelerated states). To further ensure the robustness of our findings, we restricted our final classification exclusively to subsets that exhibited consistent directional trends across both independent datasets (AIDA and OneK1K). The remaining robust subsets were then categorized into two phenotypic profiles: putative pro-aging (expanded in accelerated or depleted in decelerated cohorts) and putative pro-youthful (depleted in accelerated or expanded in decelerated cohorts).

### Definition and analysis of intra-cell-subset DEGs

The DEGs of cell subsets, including CD8^+^ naïve, MAIT, dnT, and HSPC were obtained in scRNA-seq data using the FindMarkers function in the Seurat package by comparing the top 5% contributing cells to the bottom 25% contributing cells within each cell subset, with min.pct = 0.1, logfc.threshold = 0.25 and other default parameters. After functional enrichment analysis, the top 10 significantly enriched pathways (adjusted *P*-value < 0.05, ranked by adjusted *P*-value) of each gene set were shown (Fig. 4g and Fig. S17d).

### Analysis of the SLE cohort

#### Definition of the aging trend

As previously described, we defined accelerated and decelerated aging based on predicted age residuals. To establish a healthy baseline trajectory, we applied LOWESS regression to the healthy control cohort, modeling the relationship between chronological and predicted age.

For each individual, the age residual was then calculated as the difference between their actual predicted age and the expected value derived from this healthy reference curve.

#### Trajectory analysis

Trajectory analysis was performed on four classical and two non-classical monocyte subsets. To ensure computational efficiency, a random sample of 10,000 cells from the total monocyte population was utilized as the input. Trajectory inference was carried out following the standard pipeline of the monocle R package, where dimensionality reduction was conducted using the reduceDimension function with the following parameters: max_components = 2, method = ‘DDRTree’, ncenter = 100, and lambda = 300000.

#### Identification of dynamic gene expression patterns

To characterize dynamic gene expression patterns along the evolutionary trajectory, we employed a Gaussian kernel smoothing algorithm to map discrete gene expression profiles at the single-cell level onto a continuous pseudo-time series. Specifically, we divided the total pseudo-time interval into 500 uniformly spaced reference points. For each reference point, the gene expression value was derived by calculating the weighted average of the original expression levels across all cells. The weights were determined by a Gaussian kernel function based on the distance between the cell’s actual pseudo-time and the reference point. In this process, to achieve an optimal balance between eliminating sparsity-induced noise and preserving local expression dynamics, the bandwidth of the Gaussian kernel (*σ*) was strictly set to 1% of the total pseudo-time span (sigma_ratio = 0.01).

The continuous mapping of cell identities along the pseudo-time trajectory was achieved via a combination of one-hot encoding and kernel density estimation. The original cell type labels were one-hot encoded and subjected to Gaussian weighting using the same weight matrix calculated for gene expression. A weighted score matrix for every cell type state was computed across the 500 uniformly distributed pseudo-time grid points. Subsequently, a row-wise argmax operation was applied to extract the cell category with the highest probability score at each grid point, thereby effectively defining the dominant cell state at various stages of the trajectory.

## Declarations

### Ethics approval and consent to participate

Not applicable.

### Consent for publication

Not applicable.

### Availability of data and materials

The source code for IDEAL-Age is publicly available at https://github.com/Lzcstan/IDEAL-Age under the MIT License.

All single-cell datasets analyzed in the current study are publicly available and can be downloaded from their public repositories. Specifically, the processed scRNA-seq data of the AIDA dataset can be downloaded from https://cellxgene.cziscience.com/collections/ced320a1-29f3-47c1-a735-513c7084d508 (AIDA Phase 1 Data Freeze v2); The OneK1K data can be downloaded from https://cellxgene.cziscience.com/collections/dde06e0f-ab3b-46be-96a2-a8082383c4a1; the HCA data can be downloaded from https://cellxgene.cziscience.com/collections/854c0855-23ad-4362-8b77-6b1639e7a9fc (Blood); the Sound Life data can be downloaded from https://cellxgene.cziscience.com/collections/e9360edf-b0b7-4e01-bce8-e596814f13e7; the CIMA data can be downloaded from https://db.cngb.org/trueblood/cima/resource (RNA); the siAge data can be downloaded from https://www.synapse.org/Synapse:syn61609846; the SC2018 data can be downloaded from http://gerg.gsc.riken.jp/SC2018; and the systemic lupus erythematosus data can be downloaded from https://cellxgene.cziscience.com/collections/436154da-bcf1-4130-9c8b-120ff9a888f2.

## Authors’ contributions

Y.X., D.H., Y.L., and H.W. conceived the study. D.H., Y.L., and H.W. provided overall supervision of the study. Z.L. and Y.L. designed and developed the model. F.Z., K.H., and Y.X. conducted benchmark analyses. Y.X. performed bioinformatics analyses for the gene- and cell-level interpretation interfaces. K.H. performed bioinformatics analysis for the systemic lupus erythematosus dataset under the guidance of Y.Z.. Y.X., K.H., and F.Z. wrote the main manuscript text and prepared the figures. Z.L. wrote the methodology part of the manuscript. All authors reviewed the manuscript.

## Acknowledgements

We appreciate the user-friendly data provided by the CELLxGENE database and the Human Cell Atlas community.

## Figures, tables and additional files

Additional file 1. Supplementary figures. This file contains Figs. S1-S20.

Additional file 2. Supplementary Tables. This file contains Tables. S1-S8.

