## Supplementary Figures 1-20 for "IDEAL-Age: an interpretable deep learning framework for single-cell resolution profiling of immunological aging"

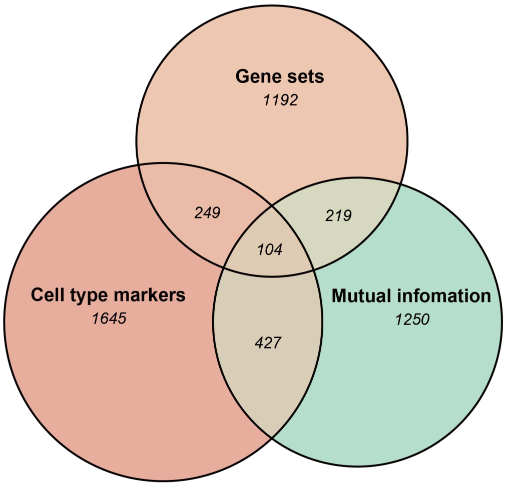


**Fig. S1 Venn plot displaying the features for the model input after selection.**

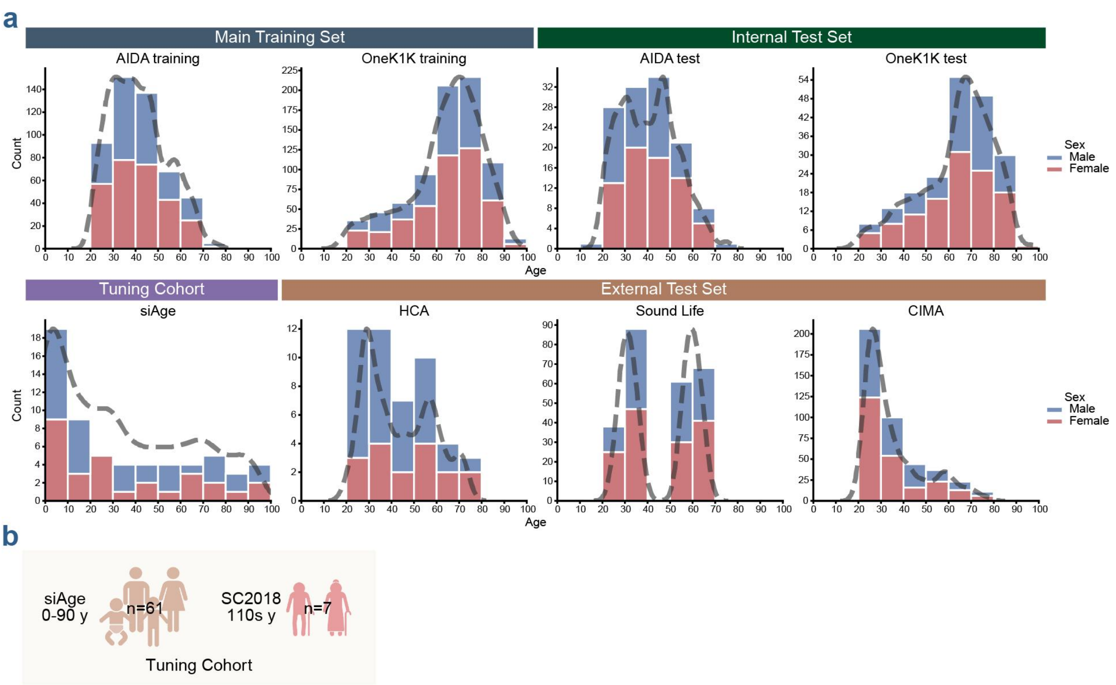


**Fig. S2 Metadata and age distribution of including cohorts. a** Age distribution of samples across individual datasets. Stacked histograms represent the frequency of samples per age decile, color-coded by biological sex (blue: male; red: female). Dashed lines indicate the kernel density estimation (KDE) of the age distribution for each cohort. **b** Graphic overview of datasets utilized for the tuning cohort of IDEAL-Age.

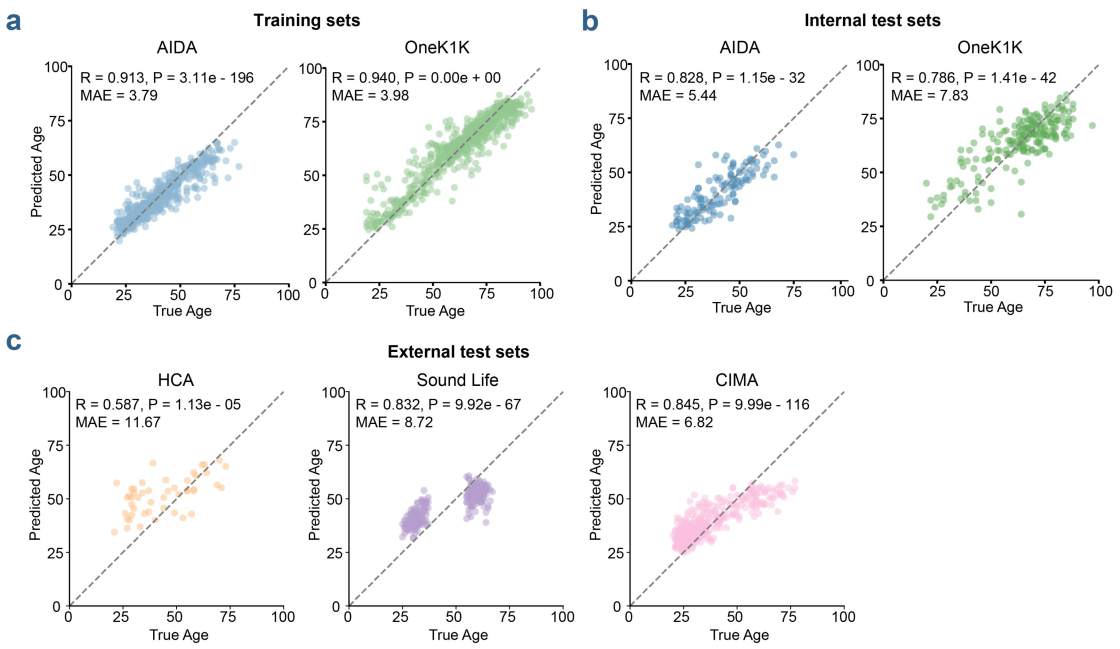


**Fig. S3 Detailed performance of IDEAL-Age on internal and external datasets.** **a-c** Scatter plots showing predicted age versus chronological age across the training (**a**), internal test (**b**) and external test (**c**) datasets of IDEAL-Age.

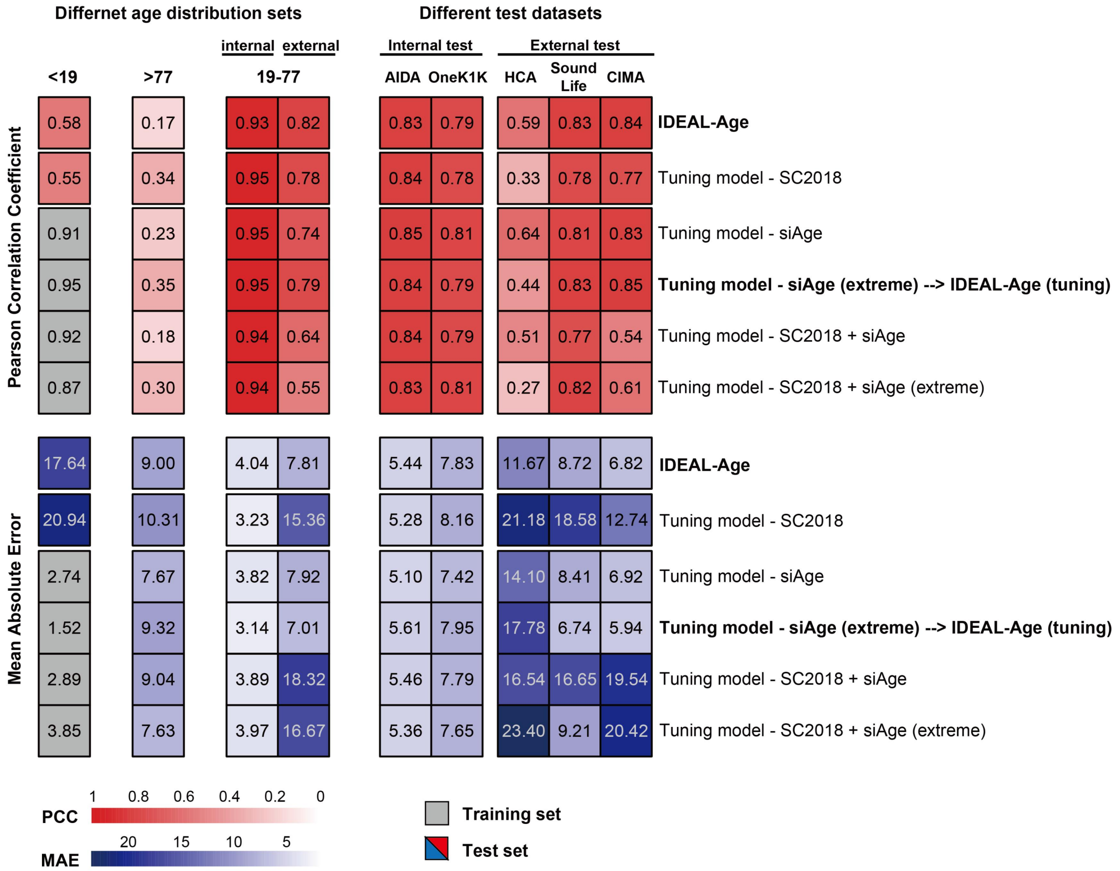


**Fig. S4 Predictive performance across various fine-tuning strategies and age distributions.** Heatmaps displaying the PCC (top) and MAE (bottom) for IDEAL-Age and five distinct model tuning configurations (rows). Performance is stratified by age distribution subsets (<19, >77, and the central 19-77 years; left panels) and across five specific transcriptomic datasets (internal tests: AIDA and OneK1K; external tests: HCA, Sound Life, and CIMA; right panels). The color intensity of each cell corresponds to the metric value, as indicated by color scales at the bottom. Note that due to cohort availability, some evaluations for the <19 age group were performed on the training set (grey boxes), whereas all other columns represent independent test sets.

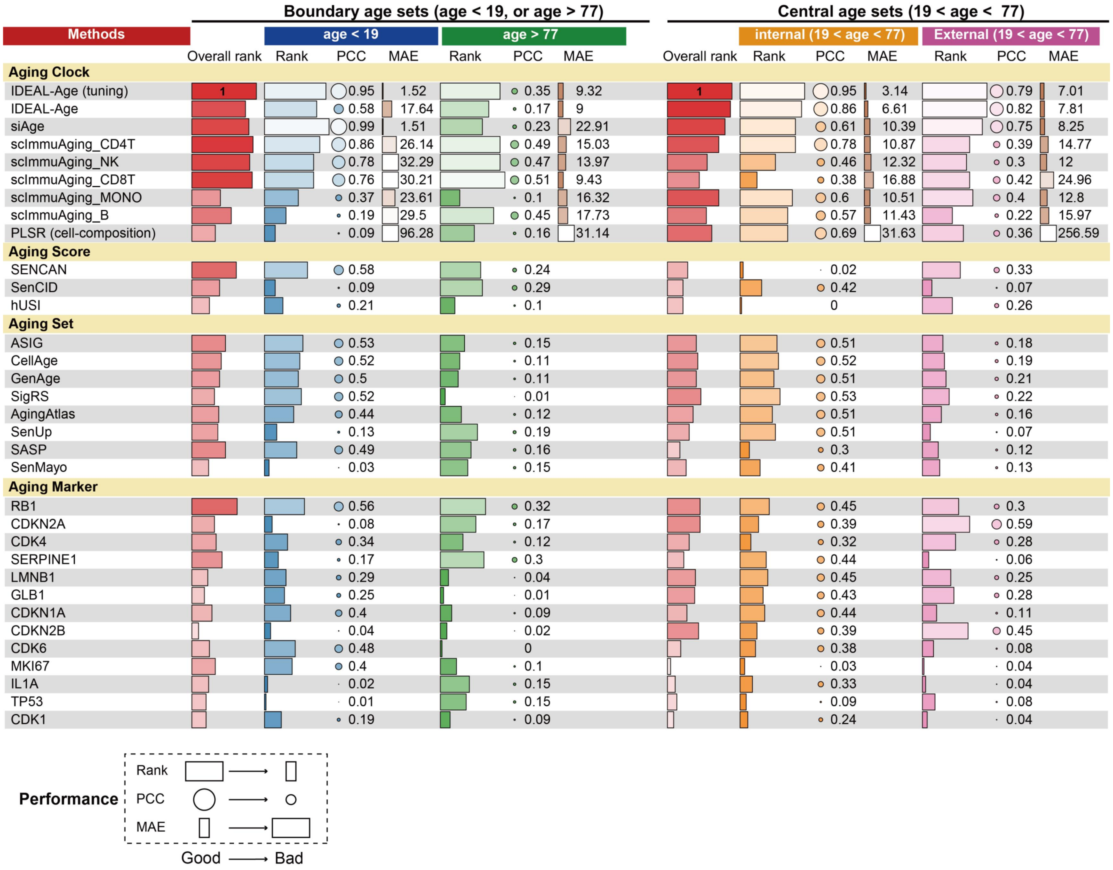


**Fig. S5 Comprehensive benchmarking of baseline and fine-tuned models across boundary and central age sets.** Heatmap summarizing predictive performance for comparing the fine-tuned model (IDEAL-Age (tuning)) against baseline and other existing models (rows). Models are categorized by their underlying evaluation strategies. Performance is evaluated independently for boundary age sets (age < 19 and age > 77; left panel) and central age sets (19 < age < 77, subdivided into internal and external cohorts; right panel). The leftmost red bars in each major section display the aggregated overall rank of the methods. The legend at the bottom denotes the relationship between geometric properties and performance quality (Good to Bad).

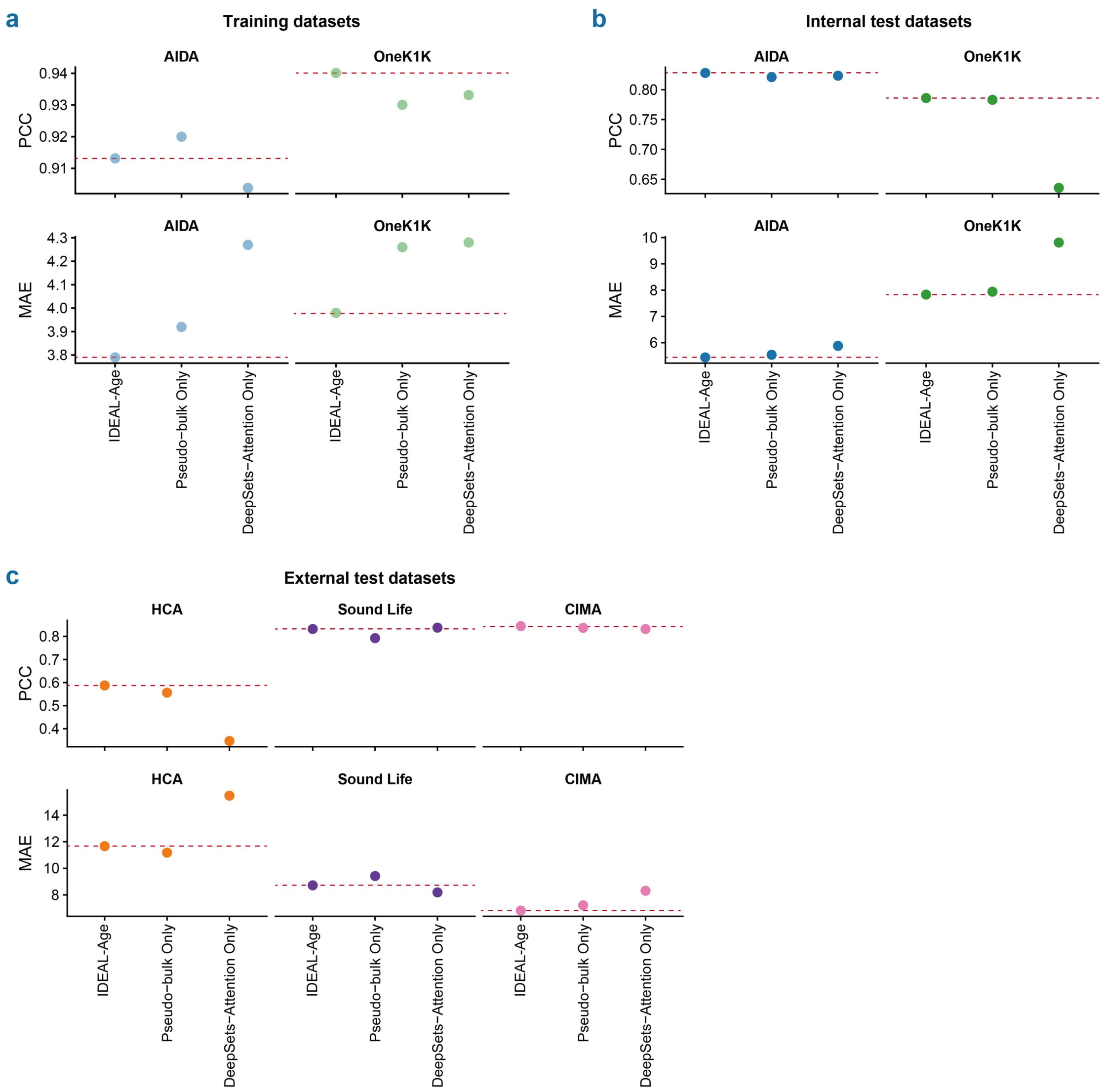


**Fig. S6 Ablation study of the IDEAL-Age architecture. a–c** Performance comparison of the full IDEAL-Age model against its ablated variants (Pseudo-bulk Only and DeepSets-Attention Only) on (**a**) training sets (AIDA, OneK1K), (**b**) internal test sets (AIDA, OneK1K), and (**c**) external test sets (HCA, Sound Life, CIMA). For each panel, the top row displays the PCC, and the bottom row displays the MAE. Red dotted lines indicate the reference performance level of the complete IDEAL-Age model.

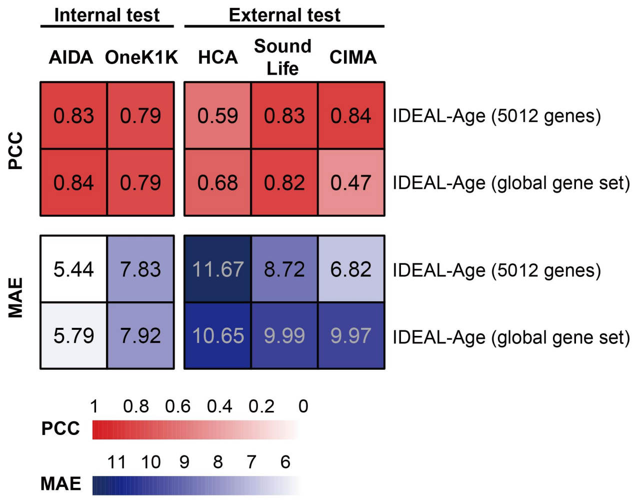


**Fig. S7 Ablation study of IDEAL-Age feature selection.** Heatmaps displaying the PCC (top panel) and MAE (bottom panel) of the IDEAL-Age model trained on the selected feature set (5,012 genes) versus a comparative model trained on the global gene set. Predictive performance was evaluated across two internal test sets (AIDA, OneK1K) and three independent external test sets (HCA, Sound Life, and CIMA).

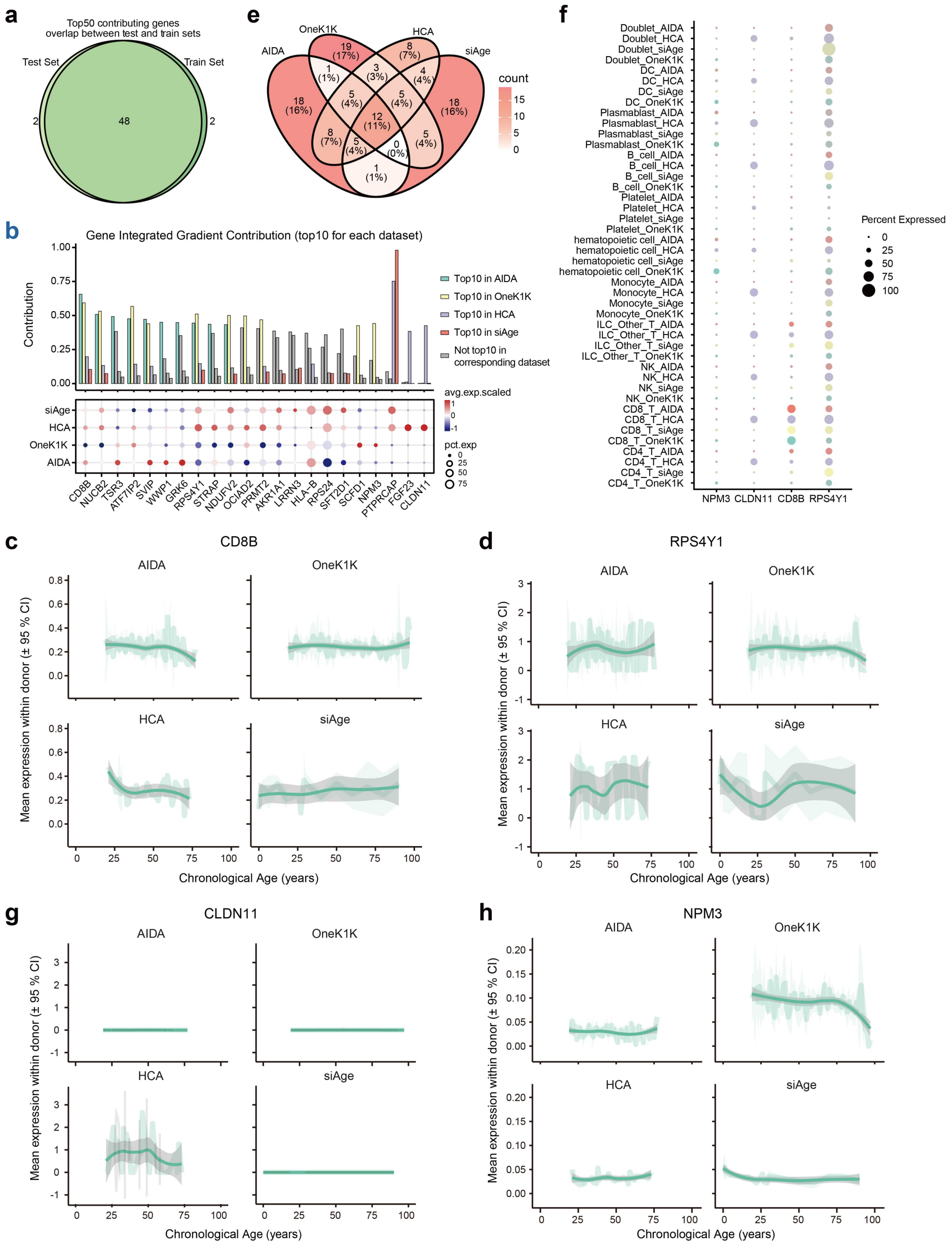

**Fig. S8 Heterogeneity of gene-level interpretability for the aging process across datasets. a** Venn diagram showing the number of overlapping genes between the top 50 contributing genes of the paired training sets and test sets. **b** Top, barplot displaying the contribution from the union of the top 10 contributing genes for each dataset. Different colors indicate respective datasets, while grey denotes genes that are not among the top 10 contributors in the corresponding dataset. Bottom, dotplot showing the expression of these genes in four datasets. **c-d** Locally Weighted Scatterplot Smoothing (LOWESS) curves showing the expression levels of *CD8B* and

*RPS4Y1* along the age trajectory in four datasets. **e** Venn diagram showing the number of overlapping genes among the top 10 contributing genes across four datasets. **f** Dot plot displaying the expression of *NPM3*, *CLDN11*, *CD8B*, and *RPS4Y1* across datasets and cell types. **g-h** LOWESS curves showing the expression levels of *NPM3* and *CLDN11* along the age trajectory in four datasets.

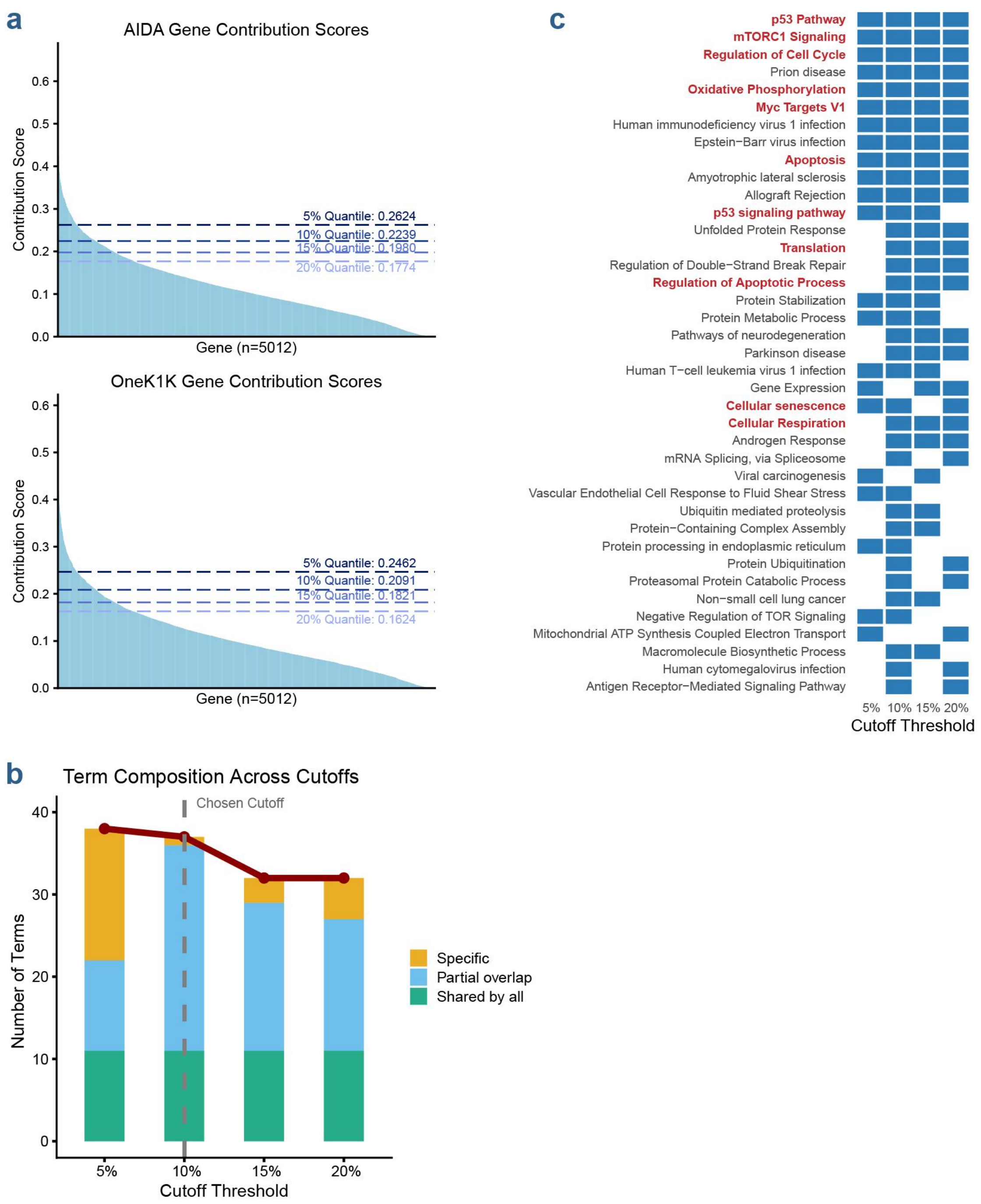


**Fig. S9 Justification and robustness analysis of gene contribution score thresholds. a** Distribution of gene contribution scores for the AIDA test set (top) and OneK1K test set (bottom). Each column represents an individual gene, ranked by its contribution score. **b** Stacked bar chart illustrating the number of enriched functional terms across four thresholds ($P < 0.05$). The colors represent unique and shared terms among the different threshold groups. **c** Heatmap displaying selected or all signaling pathways annotated under the four specified thresholds.

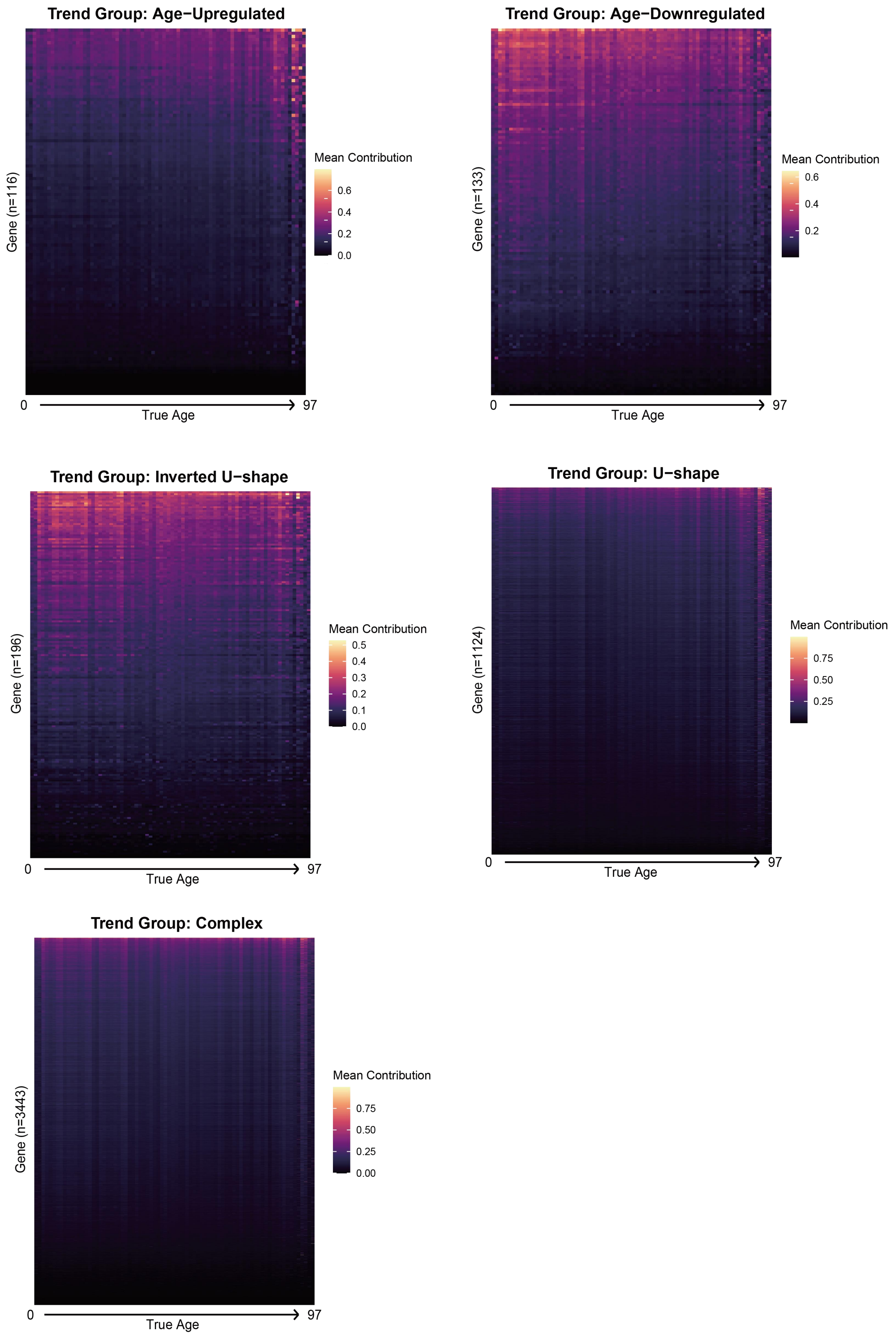


**Fig. S10** Heatmaps showing the gene contribution dynamics across the aging process in five trend groups: age-upregulated, age-downregulated, inverted U-shape, U-shape, and complex. The number of genes in each group was labeled on the y-axis.

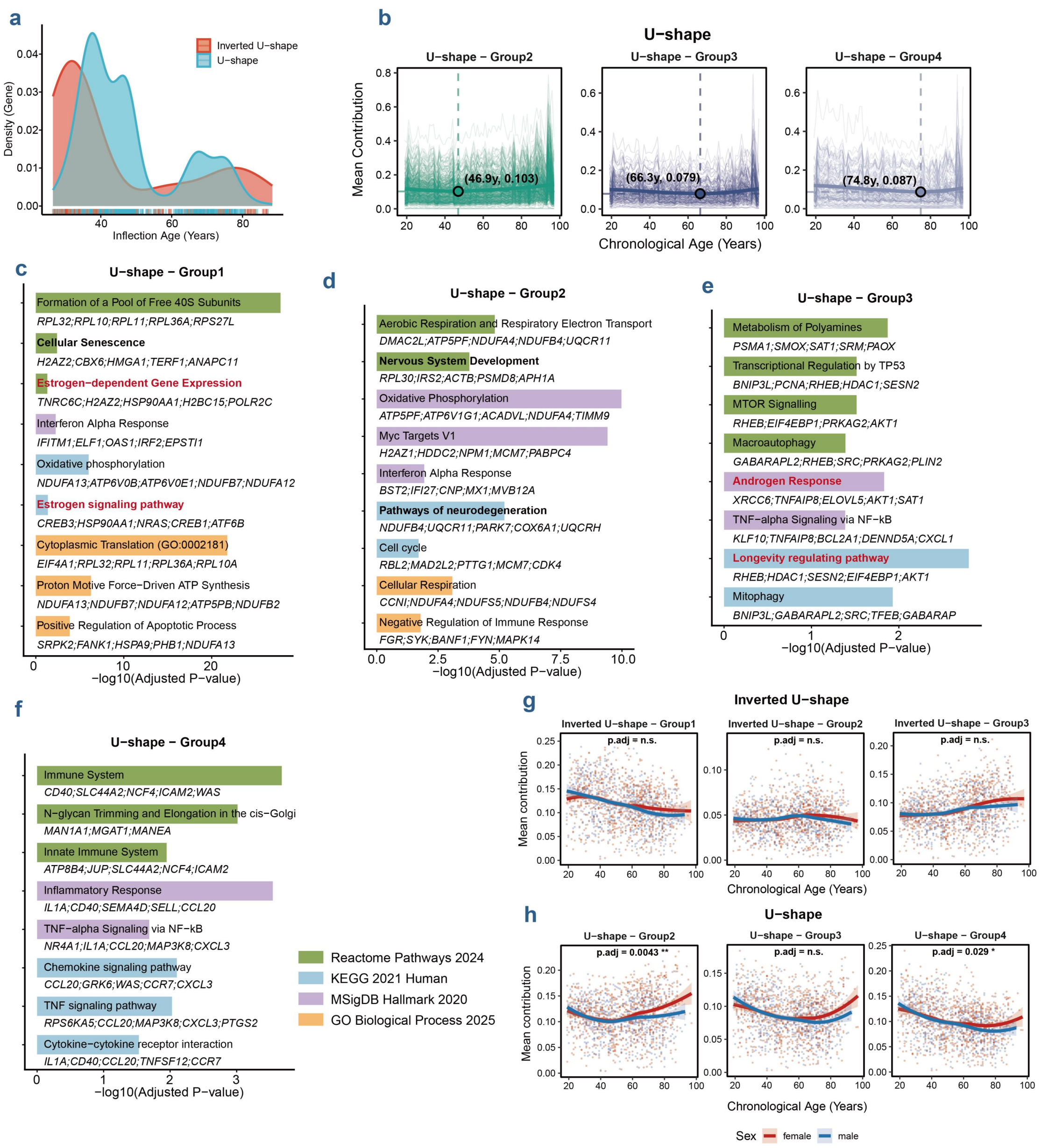


**Fig. S11 Functional enrichment and sex-specific trajectory dynamics of age-dependent gene modules. a** Density plot illustrating the overall distribution of inflection ages for genes exhibiting U-shape (blue) and Inverted U-shape (red) trajectories. Rug marks at the bottom represent individual gene inflection points. **b** Longitudinal trajectories of mean contribution across chronological age for U-shape group 2 to 4. Faded thin lines represent the trajectories of individual genes, while the bold solid lines represent the LOWESS average trajectory for each respective cluster. Dashed lines and colored circles indicate the exact group-level inflection point (nadir for U-shape group), with the precise age and mean contribution coordinates labeled. **c-f** Pathway enrichment analysis for the U-shape groups (U1-U4). Bar lengths represent -log$_{10}$(Adjusted *P*-value) for selected significantly enriched pathways (adjusted $P < 0.05$). **g-h** Sex-stratified mean contributions across chronological age for Inverted U-shape (**g**) and U-shape groups (**h**). Dots represent individual samples (females in red, males in blue). Solid lines depict LOWESS with shaded areas representing 95% confidence intervals. Overall sex differences across the lifespan were assessed using a likelihood ratio test comparing nested Generalized Additive Models (GAMs). Resulting *P*-values were corrected for multiple testing using the Benjamini-Hochberg method (p.adj). $*P_{adj} < 0.05$, $**P_{adj} < 0.01$, $***P_{adj} < 0.001$; n.s., not significant.

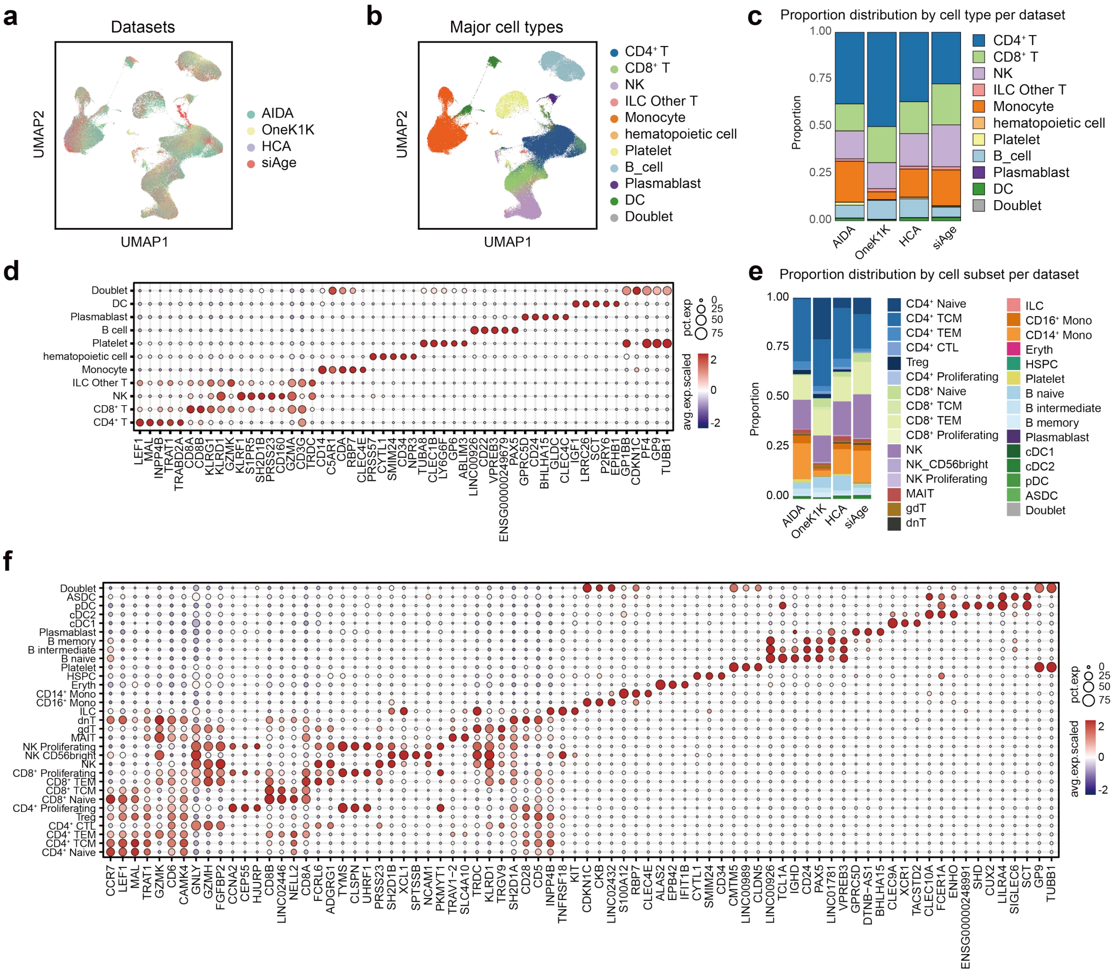


**Fig. S12** Integrated scRNA-seq data annotation. **a** UMAP plot showing the cell distribution of their source datasets. **b** UMAP plot showing 11 major cell types identified in the integrated scRNA-seq data. **c** Stacked bar plot showing the major cell type composition of 4 datasets. **d** Dot plot showing the expression of top 5 specific marker genes for each major cell type in the integrated scRNA-seq data. **e** Stacked bar plot showing the fine-grained cell subset composition of 4 datasets. **f** Dot plot showing the expression of top 3 specific markers for each fine-grained cell subset in the integrated scRNA-seq data.

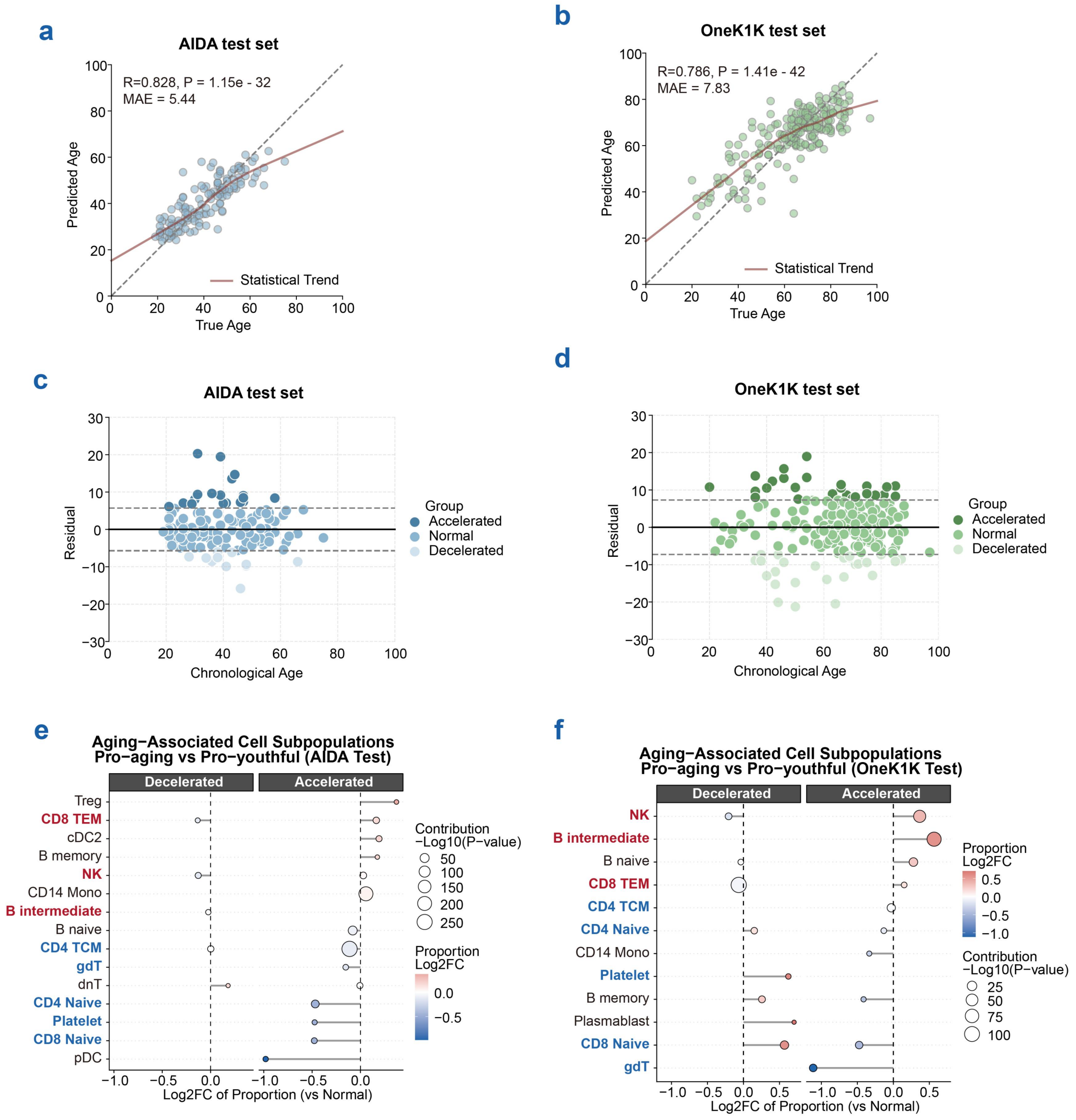


**Fig. S13 Identification of aging-associated cell subpopulations based on predicted aging trajectories. a-b** Scatter plots showing the correlation between chronological age and predicted age within the AIDA (**a**) and OneK1K (**b**) test sets. Solid red lines represent LOWESS regression fits, and dashed grey lines indicate the identity line (y = x). **c-d** Distribution of prediction residuals across chronological age. Donors are stratified into three groups: accelerated aging (residual > 1 standard deviation [s.d.]), normal aging (within ±1 s.d.), and decelerated aging (residual < -1 s.d.). Dashed horizontal lines denote the ±1 s.d. thresholds, and solid black lines indicate a residual of 0. **e-f** Shifts in relative proportions of cell subpopulations across decelerated and accelerated cohorts in the AIDA (**e**) and OneK1K (**f**) test sets. The x-axis indicates the log2 fold change (Log2FC) of cell proportions relative to the normal aging cohort. Dot size reflects the statistical significance of the predictive contribution score, calculated via the Wilcoxon rank-sum test. Cell types labeled in red or blue on the y-axis denote putative pro-aging or pro-youthful subpopulations, respectively.

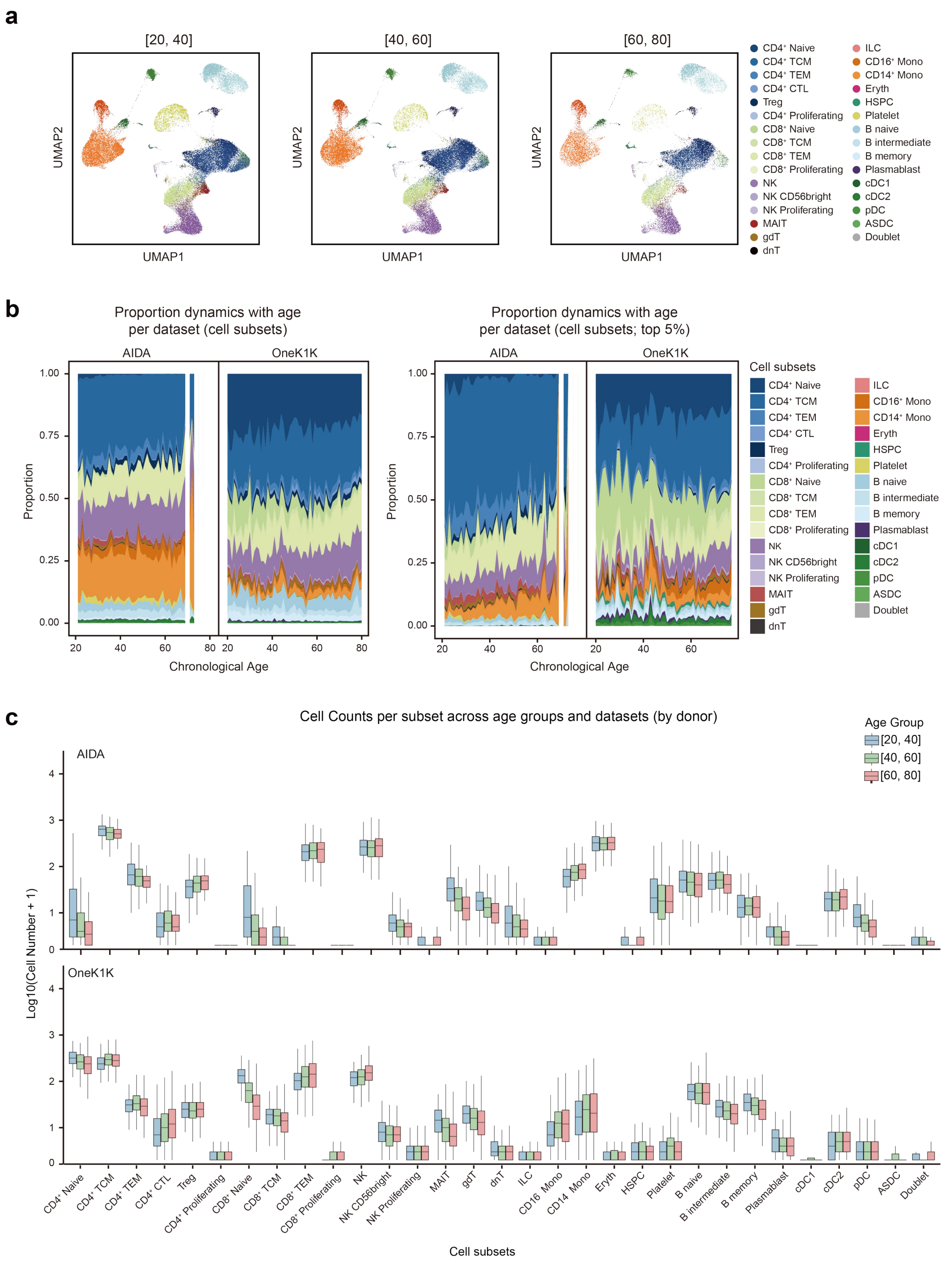


**Fig. S14** Characteristics of the dataset combining from AIDA and OneK1K with age from 20 to 80. **a** UMAP plots displaying the cell subsets distribution of each age group. **b** Stacked area plots displaying the proportional dynamics of cell subsets of total cells (left) and the top 5% contributing cells (right) across age for each dataset. **c** Boxplots showing the distribution of cell numbers per donor for each cell subset across age groups within each dataset. Boxes represent the median and interquartile range (IQR), with whiskers extending to 1.5× IQR.

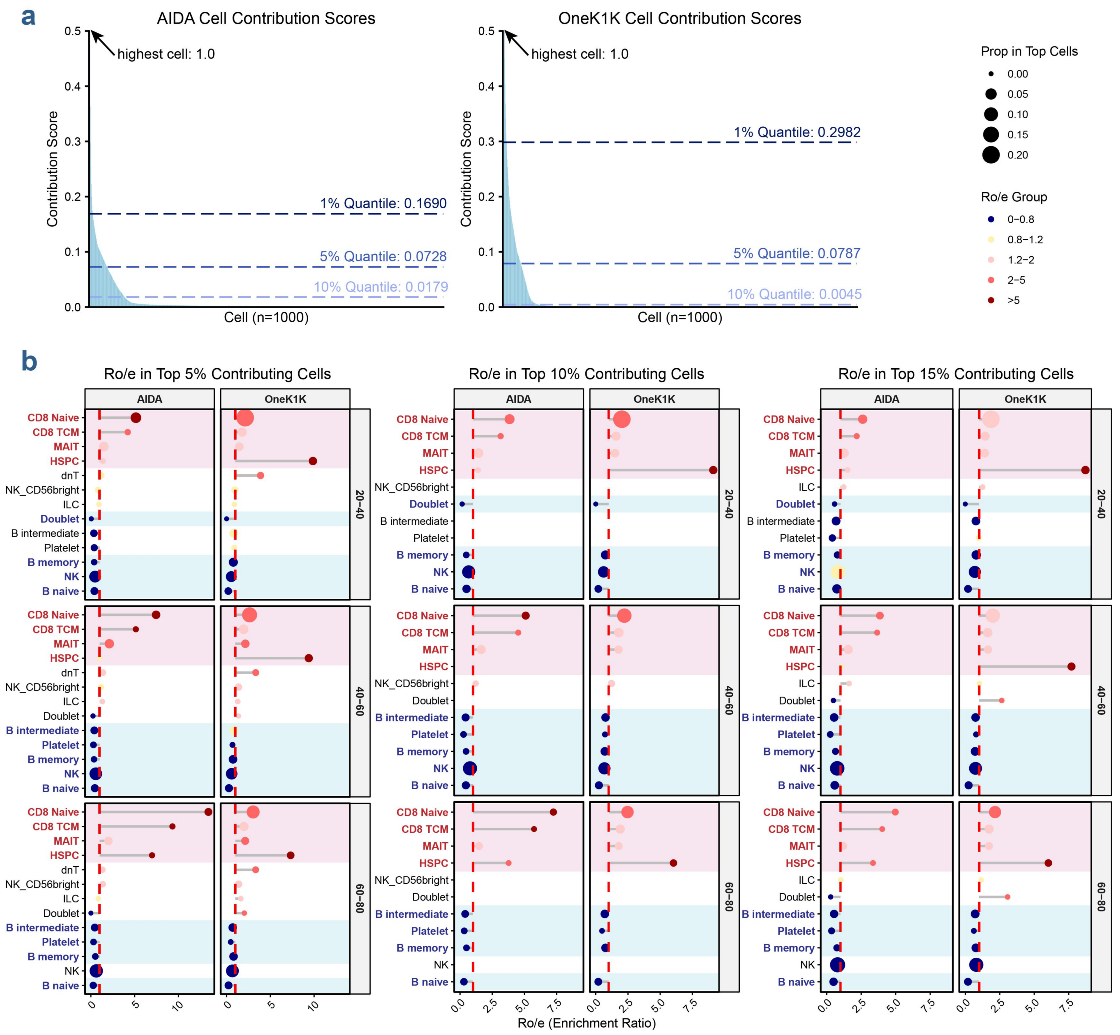


**Fig. S15 Justification and robustness analysis of cell contribution score thresholds. a** Distribution of cell contribution scores for the AIDA test set (left) and OneK1K test set (right). Each column represents an individual cell, ranked by its contribution score. For visualization purposes, 1,000 cells were uniformly sampled across the entire population based on their ranked scores. **b** Lollipop plot showing the relative enrichment (Ro/e) of cell subsets across age groups and datasets at three thresholds. Dot size is proportional to the relative abundance of the top 5% contributing cells, and color indicates Ro/e groups. Labels and background in red or blue denote cell subsets with consistent high or low enrichment trends across thresholds. The red dashed line marks the baseline Ro/e = 1.

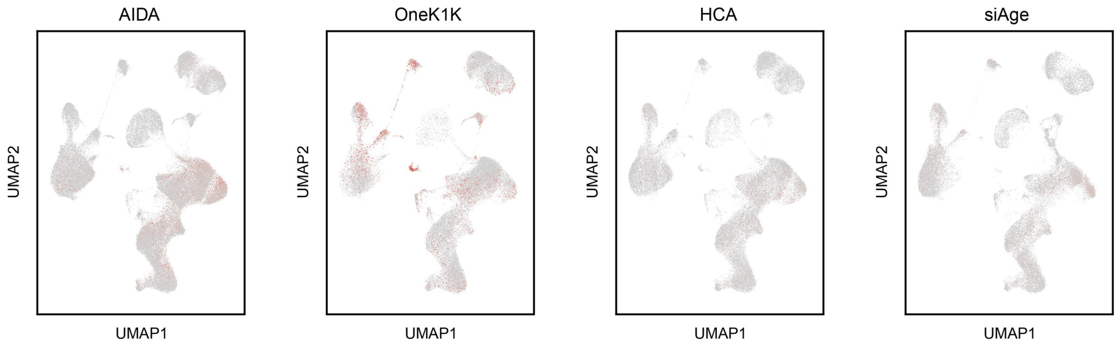


**Fig. S16** UMAP plot displaying the cell-level contribution in each dataset.

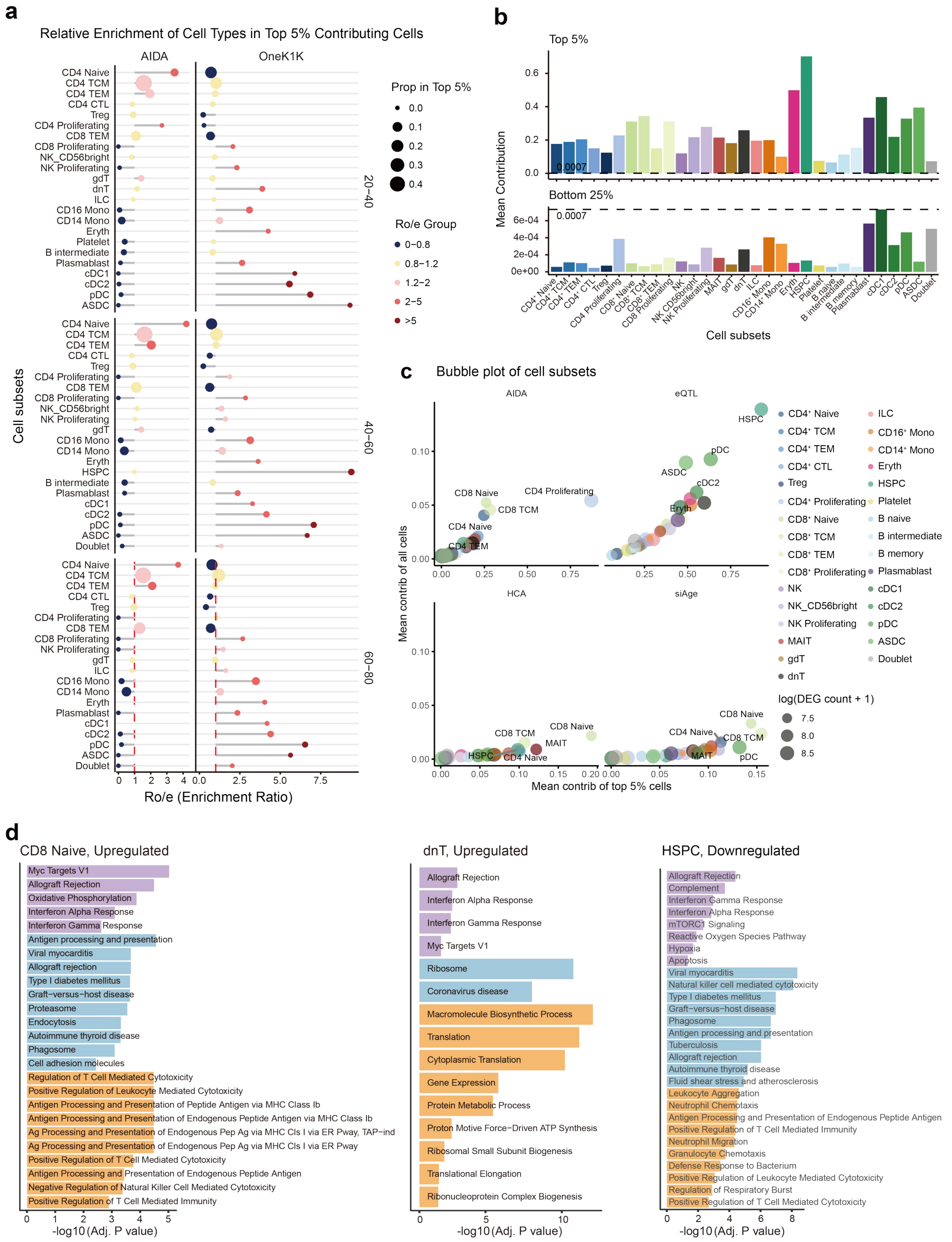

**Fig. S17** Comparison of top 5% and bottom 25% contributing cells in each cell subset. **a** Lollipop plot showing the relative enrichment (Ro/e) of cell subsets across age groups and datasets. Dot size is proportional to the relative abundance of the top 5% contributing cells, and color indicating Ro/e groups. The red dashed line marks the baseline Ro/e = 1. **b** Bar plots showing the mean contribution scores of top 5% cells (top) and bottom 25% cells (bottom) in each cell subset. The dashed lines represent the maximum contribution score thresholds of the bottom 25% cell group. **c** Bubble plots displaying the mean contribution scores of each cell subset (y-axis), and the mean contribution of the top 5% cells within certain cell subsets (x-axis), split by dataset. Colors represent different cell subsets. **d** Bar plots depicting functional enrichment terms of genes upregulated in CD8$^+$ naïve T and dnT, and genes downregulated in HSPC, comparing the top 5% and bottom 25% contributing cells.

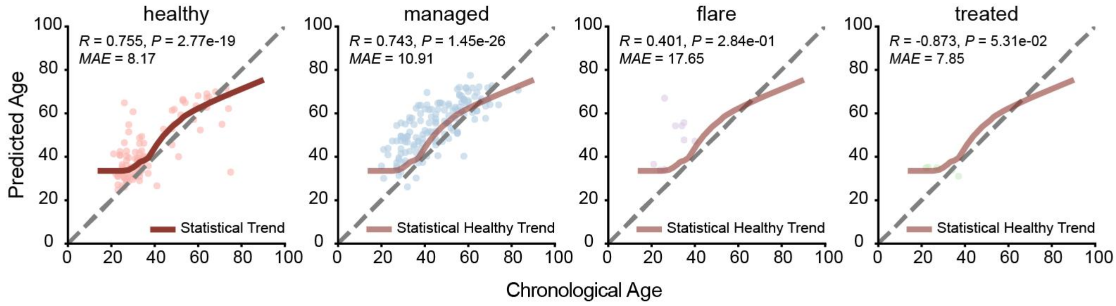


**Fig. S18 Regression analysis of aging trends based on healthy control samples.** Scatter plot illustrating the correlation between predicted age and chronological age. The statistical trend for healthy controls and the "statistical healthy trend" for SLE patients were derived using LOWESS regression fitted to the healthy cohort.

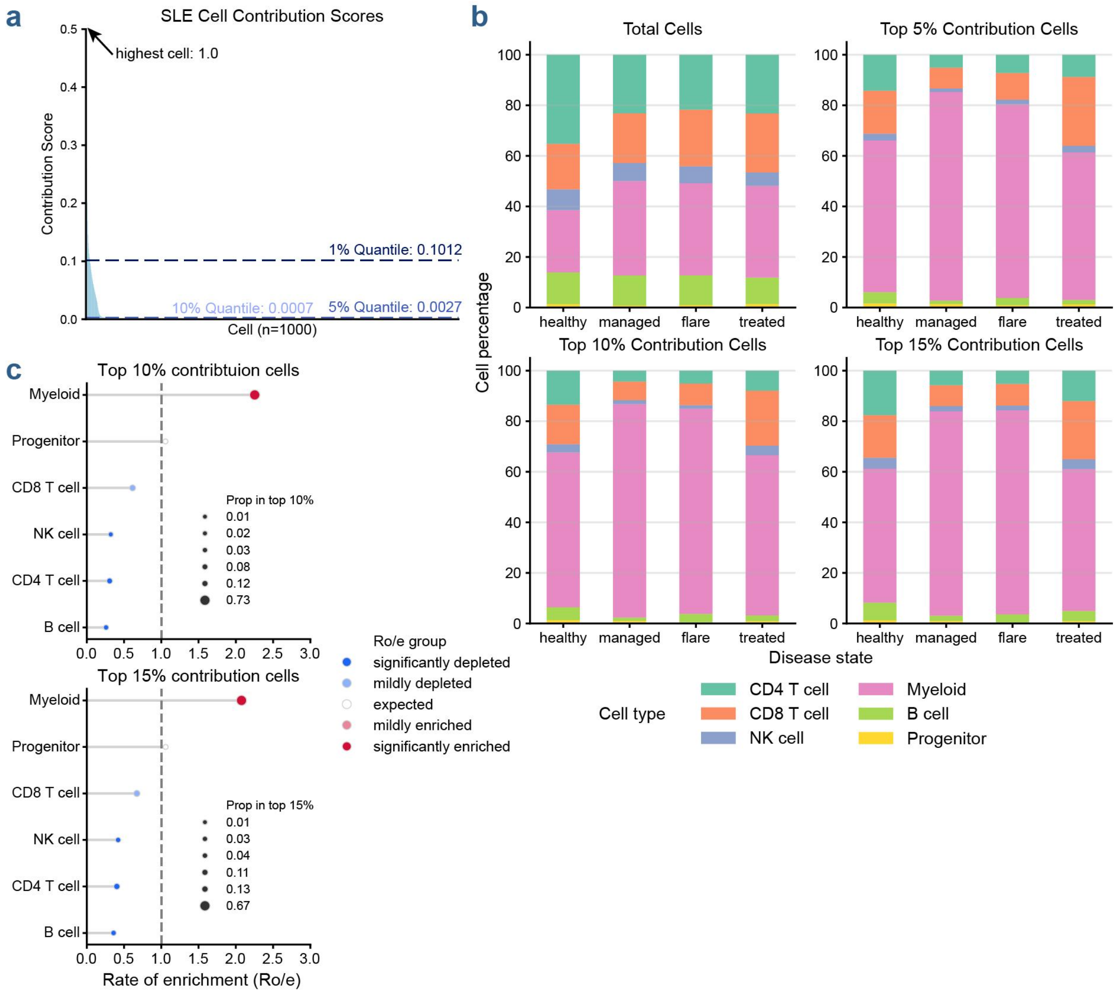


**Fig. S19 Robustness validation of cell filtering and selection in the SLE cohort. a** Distribution of cell contribution scores. Each column represents an individual cell, ranked by its contribution score. For visualization purposes, 1,000 cells were uniformly sampled from the entire population based on their ranked scores. **b** Stacked bar plots of cell type composition across varying thresholds of contribution scores (total cells, top 5%, 10%, and 15%). **c** Lollipop plot illustrating the enrichment rates of specific cell types under different filtering thresholds. Dot size is proportional to the relative abundance of the top contributing cells, and color indicates Ro/e groups. The gray dashed line marks the baseline Ro/e = 1.

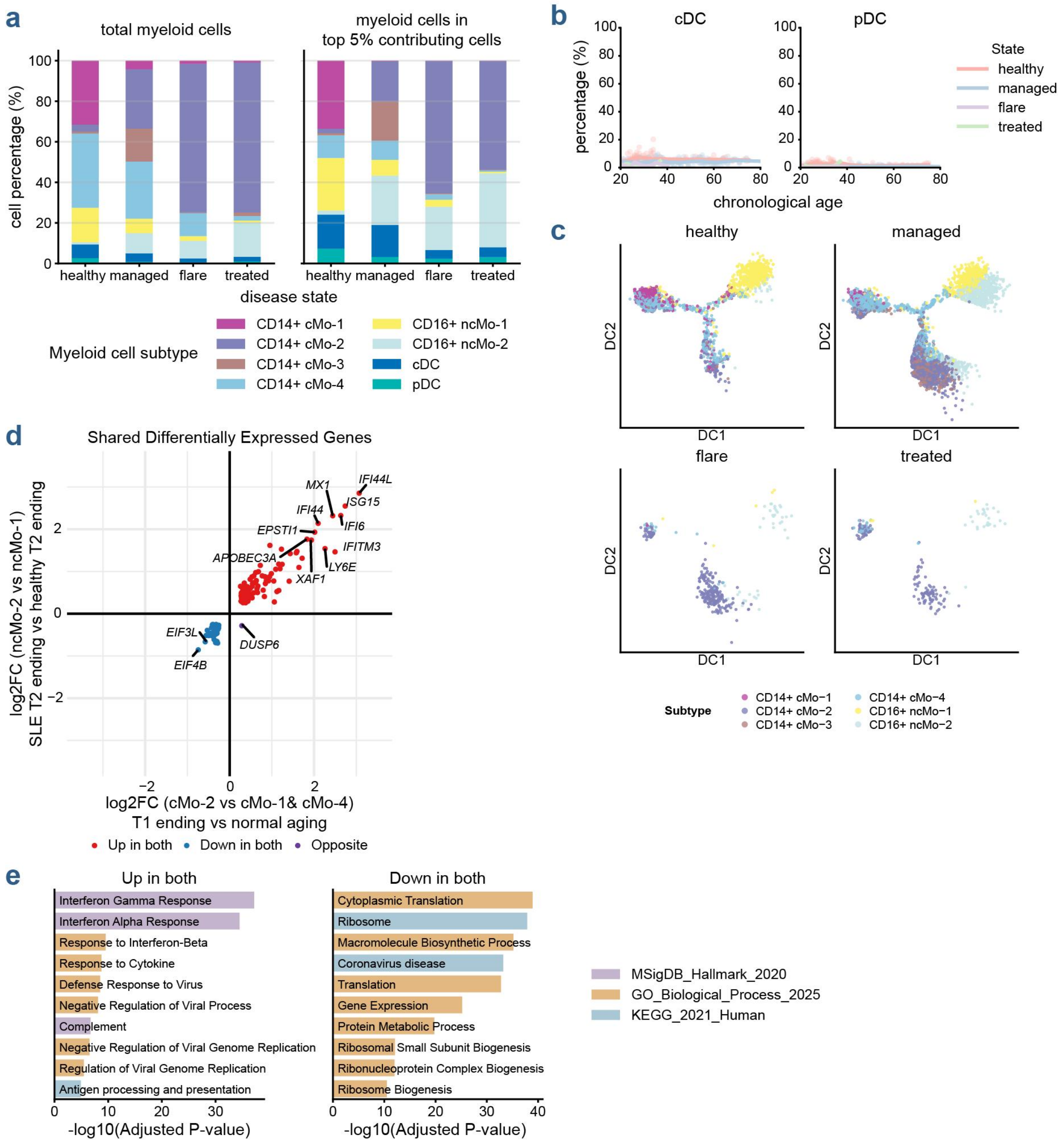


**Fig. S20 Transcriptional dynamics and trajectory analysis of myeloid cell subsets across disease states. a** Stacked bar charts showing the proportional distribution of myeloid cell subsets. The left panel displays the composition within total myeloid cell population across different clinical states (healthy, managed, flare, and treated). The right panel shows the distribution specifically within the top 5% of cells with the highest contribution scores. **b** Scatter plots illustrating the percentage of conventional dendritic cells (cDCs) and plasmacytoid dendritic cells (pDCs) relative to total myeloid cells across chronological age in different clinical states. **c** Diffusion map embeddings (DC1 vs DC2) revealing the developmental trajectories of myeloid subsets, stratified by disease state. Single cells are colored according to their specific subpopulation annotations. **d** Scatter plot comparing the $\log_2$ fold change of shared differentially expressed genes (DEGs) at the SLE T2 ending (ncMo-2 vs. ncMo-1) versus the T1 ending (cMo-2 vs. cMo-1/4). Genes that are highlighted as consistently upregulated (red), consistently downregulated (blue), or exhibit opposite expression trends (purple) across the two trajectories. **e** Functional enrichment analysis of the shared upregulated (left) and downregulated (right) genes identified in **d**, utilizing MSigDB Hallmark, GO Biological Process, and KEGG databases. The x-axis represents enrichment significance ($-\log_{10}$(Adjusted *P*-value)).
